# A retinal adrenergic module tunes mammalian visual evolution

**DOI:** 10.64898/2026.08.15.744996

**Authors:** Fu-Sheng Tang, Miao-Miao Kong, Ya-Ru Luo, Zi-Xuan Wang, Min Gao, Lin-Jie Rao, Jun-Bin Liu, Ting-Ting Zhang, Si-Yun Chen, Yun Cheng, Bin Gou, Cheng Yang, Hong-Bo Yu, Jing-Tao Lilue, Wei Li, Jiang-Bin Ke

**Author notes:** **Correspondence and requests for materials** should be addressed to Jiang-Bin Ke. These authors contributed equally (Fu-Sheng Tang, Miao-Miao Kong).

## Abstract

How conserved neural circuits are modified during mammalian evolution remains poorly understood. Here we combine cross-species single-cell transcriptomics, in situ validation, retinal physiology, and conditional genetics to identify a superorder-associated adrenergic module in the mammalian retina. We find that *ADRB1*, which encodes the β_1_-adrenergic receptor, is uniquely expressed in rod bipolar cells of sampled Euarchontoglires, but is absent from homologous cells in sampled Laurasiatheria and Marsupialia. In mice, β_1_-adrenergic receptor localizes to rod bipolar cell terminals and boosts transmission to AII amacrine cells through G_s_-adenylyl cyclase-cAMP-PKA signaling pathway. This modulation enhances synchronous release, accelerates downstream ganglion cell output, and increases scotopic electroretinographic responses, while rod-bipolar-cell-specific *Adrb1* deletion abolishes norepinephrine-induced enhancement without disrupting baseline vision. In the diurnal tree shrew, a Euarchontoglires species with a cone-dominated retina, *ADRB1* is instead redeployed from rod bipolar cells to cone photoreceptors. These findings reveal an evolutionarily mobile neuromodulatory module that tunes retinal computation according to visual ecology.

## Introduction

The mammalian retina contains a conserved set of neuronal classes that transform photoreceptor signals into parallel output channels for vision^1–5^. This conservation makes the retina a powerful system for asking how evolution modifies neural circuits without rebuilding them from first principles^3,5,6^. Comparative genomics has clarified the broad relationships among placental mammals, including the ancient divergence between Euarchontoglires and Laurasiatheria^7–9^, but how such large-scale evolutionary splits are reflected in cell-type-specific circuit function remains poorly understood.

Sensory evolution is often considered in terms of changes in receptors, sensory organs or behavior^10–15^. In vision, classic examples include opsin evolution, trichromacy, nocturnality, and specialization of photoreceptor composition^3,5,10,12,14,16–23^. Yet adaptive changes may also occur at synaptic and modulatory nodes within otherwise conserved circuits^3,6,18,21^. Such changes provide an efficient evolutionary strategy: by altering where a neuromodulatory receptor is expressed, the same transmitter system can be coupled to distinct computations in different species.

Rod-mediated vision is particularly suited to this type of modulation. Rod photoreceptors can detect single photons, and their signals are relayed through the rod bipolar cell pathway, in which rod bipolar cells transmit visual information to AII amacrine cells and subsequently to ON and OFF ganglion cell pathways^1,2,19,24–28^. This circuit is optimized for sensitivity, but its output must also be adjusted according to behavioral state, ambient illumination, and ecological demand^21,26,27,29–31^. Neuromodulators, including catecholamines, are known to reconfigure retinal processing^21,32,33^, but whether the cellular targets of neuromodulation have themselves evolved across mammalian clades remains largely unknown.

Here we identify a retinal adrenergic module associated with Euarchontoglires. By analyzing cross-species single-cell transcriptomic datasets (Gene Expression Omnibus (GEO) accession numbers: GSE237215, GSE63472, and GSE118480; European Genome-phenome Archive: EGAS00001004561)^5,34–36^ and a newly generated rabbit retinal dataset (GEO accession number: GSE336783), we found that *ADRB1*, which encodes β_1_-adrenergic receptor (β_1_AR), is expressed in rod bipolar cells across sampled Euarchontoglires but not in homologous cells from sampled Laurasiatheria or Marsupialia retina. We then used the mouse retina to determine whether this evolutionary molecular signature has functional consequences. β_1_AR activation enhanced rod bipolar cell output, strengthened AII amacrine cell responses, increased ganglion cell signaling, and boosted scotopic electroretinographic responses in vivo. Conditional deletion of *Adrb1* from rod bipolar cells abolished this norepinephrine-dependent enhancement while sparing baseline retinal function. Finally, we found that the diurnal tree shrew, a Euarchontoglires species with a cone-dominated retina^37^, has shifted *ADRB1* expression from rod bipolar cells to cone photoreceptors. Together, these results reveal a cell-type-specific, evolutionarily redeployable neuromodulatory mechanism for tuning mammalian visual function.

## Results

### *ADRB1* marks Euarchontoglires rod bipolar cells

To search for cell-type-specific molecular features that distinguish major mammalian lineages, we analyzed retinal single-cell RNA-sequencing data from Euarchontoglires, Laurasiatheria, and Marsupialia, and integrated these data with a newly generated rabbit retinal single-cell dataset. Major retinal cell classes, including rods, cones, horizontal cells, bipolar cells, amacrine cells, ganglion cells, and Müller glia, were identified using conserved marker genes and established retinal cell-type annotations (Fig. 1a and Extended Data Fig. 1).

**Fig. 1.**
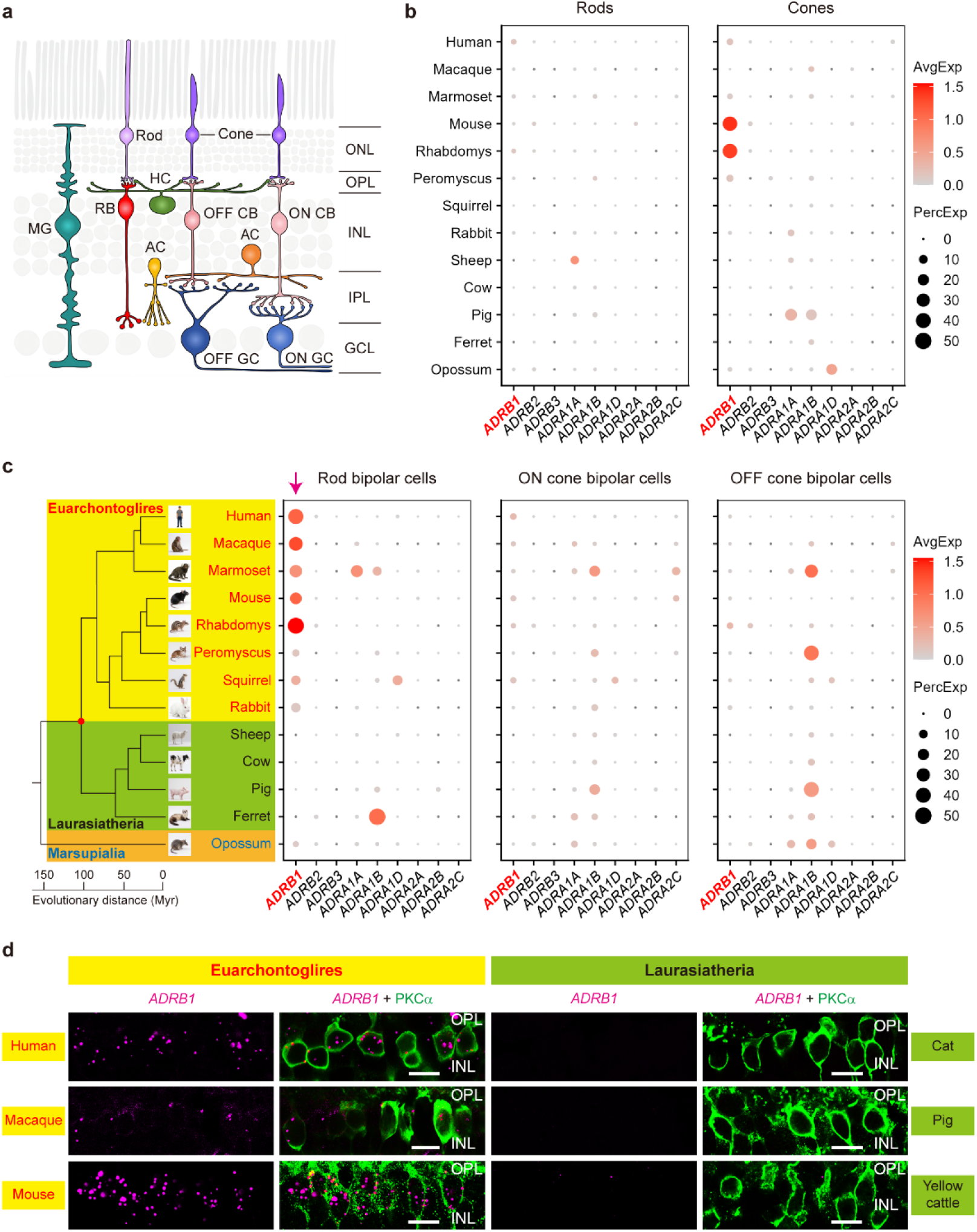
*ADRB1* expression in rod bipolar cells is associated with Euarchontoglires. **a**, Schematic of the mammalian retinal circuit, highlighting photoreceptors (rods and cones), horizontal cells (HCs), rod bipolar (RB) cells, ON and OFF cone bipolar (CB) cells, amacrine cells (ACs), ON and OFF ganglion cells (GCs), and Müller glia (MG). The outer nuclear layer (ONL), inner nuclear layer (INL), ganglion cell layer (GCL), outer plexiform layer (OPL), and inner plexiform layer (IPL) are indicated. **b**, Dot plots showing adrenergic receptor gene expression in rods and cones across 13 species. Dot size indicates the fraction of expressing cells and color indicates average normalized expression. **c**, Phylogenetic overview of the 13 mammalian species analyzed (left) and dot plots showing *ADRB1* expression in RBs of sampled Euarchontoglires, but not in sampled Laurasiatheria or Marsupialia. Arrow indicates the RB-specific Euarchontoglires-associated pattern. **d**, RNAscope validation of *ADRB1* transcripts in PKCα-positive rod bipolar cells from representative Euarchontoglires (human, macaque, mouse) and Laurasiatheria (cat, pig, yellow cattle) retinas. Biological replicates: human, macaque, cat, pig, and yellow cattle, n = 2 each; mouse, n = 3. Scale bars, 10 μm. See also Extended Data Figs. 1-3.

We first focused on adrenergic receptor genes because neuromodulatory receptors can couple behavioral state to sensory processing^38–42^. Across retinal cell types, most adrenergic receptors showed sparse or species-restricted expression (Figs. 1b,c and Extended Data Fig. 2). A striking exception was *ADRB1*. In rod bipolar cells, *ADRB1* was consistently detected across sampled Euarchontoglires, including primates, rodents, and rabbit, whereas it was absent or nearly absent from rod bipolar cells of sampled Laurasiatheria and from opossum (Fig. 1c). This pattern was not observed for other adrenergic receptor genes and was not reproduced in other major retinal cell classes (Figs. 1b,c and Extended Data Fig. 2).

Uniform manifold approximation and projection (UMAP) visualization confirmed that *ADRB1* expression in Euarchontoglires was concentrated in rod bipolar cells, with additional cone expression in a subset of species (Extended Data Fig. 1). By contrast, Laurasiatheria retinas lacked comparable *ADRB1* expression in rod bipolar cells (Extended Data Fig. 1). To validate this transcriptomic pattern at the tissue level, we performed RNAscope in situ hybridization for *ADRB1* together with PKCα, a specific marker of rod bipolar cells. *ADRB1* transcripts were detected in PKCα-positive rod bipolar cells in human, macaque, and mouse retinas, but not in cat, pig, or yellow cattle retinas (Fig. 1d). Thus, *ADRB1* expression in rod bipolar cells represents a robust, cell-type-specific molecular feature associated with Euarchontoglires.

Differential expression analysis identified additional genes enriched in rod bipolar cells or photoreceptors of either Euarchontoglires or Laurasiatheria (Extended Data Fig. 3). However, *ADRB1* was distinctive because it encodes a neuromodulatory receptor with an immediately testable role in retinal computation. We therefore next asked whether this evolutionary signature has functional consequences.

### β_1_AR modulates rod bipolar cell output

We used the mouse retina as an experimentally tractable Euarchontoglires model. Immunolabelling showed that β_1_AR protein was enriched at rod bipolar cell axon terminals in the inner plexiform layer (Figs. 2a,b), suggesting that the receptor may regulate synaptic output from rod bipolar cells to AII amacrine cells.

**Fig. 2.**
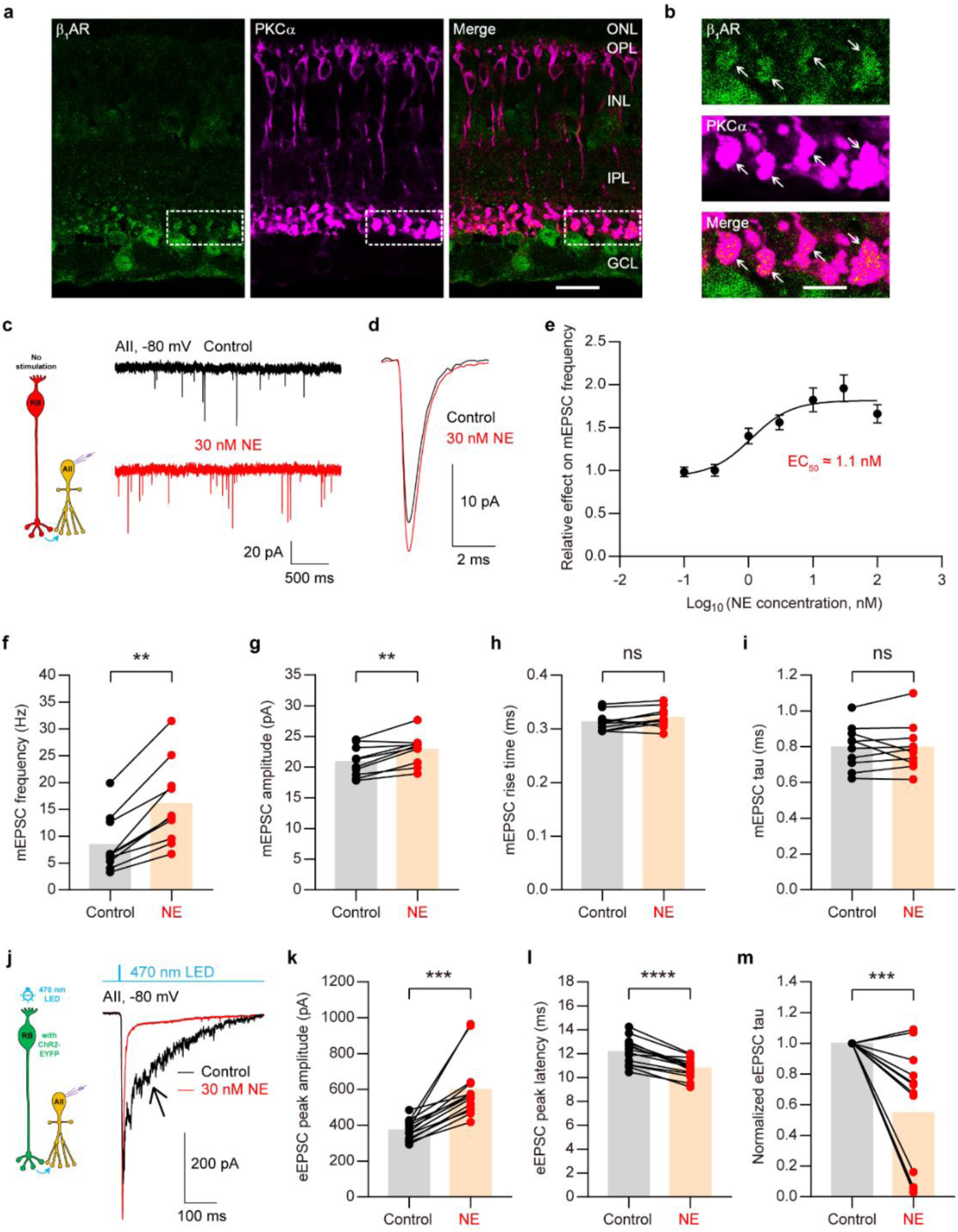
β_1_AR activation tunes rod bipolar cell synaptic release. **a**, β_1_AR immunolabelling in mouse retina showing enrichment at PKCα-positive rod bipolar cell terminals. ONL, outer nuclear layer; OPL, outer plexiform layer; INL, inner nuclear layer; IPL, inner plexiform layer; GCL, ganglion cell layer. Scale bar, 20 μm. **b**, Enlarged view of the boxed region in **a**. Arrows indicate rod bipolar cell terminals. Scale bar, 5 μm. Images in **a** and **b** are representative of n ≥ 3 experiments. **c**, Recording configuration for miniature excitatory postsynaptic currents (mEPSCs) from AII amacrine cells (left), and representative mEPSCs before and after norepinephrine (NE) application (right). **d**, Average mEPSC waveforms from the same AII in **c** under control and NE conditions. **e**, Dose-response relationship for the effect of NE on mEPSC frequency. EC_50_ ≈ 1.1 nM. Data are mean ± s.e.m. **f-i**, Quantification of mEPSC frequency (**f**; P = 0.0020), amplitude (**g**; P = 0.0012), rise time (**h**; P = 0.0569), and decay tau (**i**; P = 0.9661). n = 10 cells from 5 mice. **j**, Recording configuration for optogenetically evoked rod bipolar cell input to AII amacrine cells (left), and representative evoked excitatory postsynaptic currents (eEPSCs) showing fast and slow components before and after NE application (right). Arrow indicates the slow component. **k-m**, Quantification showing that NE enhances the fast component (**k**; P = 0.0001), shortens the time to peak (**l**; P < 0.0001), and suppresses the slow component (**m**; P = 0.0006). n = 14 cells from 7 mice. Paired *t*-test or Wilcoxon test. **P < 0.01; ***P < 0.001; ****P < 0.0001; ns, not significant. Unless otherwise indicated, data are presented as individual points with means; error bars are not shown. See also Extended Data Figs. 4-6.

We first recorded miniature excitatory postsynaptic currents from AII amacrine cells, which receive glutamatergic input from rod bipolar cells (Fig. 2c). Bath application of norepinephrine increased both the frequency and amplitude of miniature events, with an EC_50_ of approximately 1.1 nM, without altering event kinetics (Figs. 2c-i). Thus, β_1_AR-dependent modulation of the rod bipolar cell→AII synapse operates with nanomolar sensitivity, suggesting that this synapse is highly sensitive to norepinephrine. The increase in frequency indicates enhanced presynaptic release from rod bipolar cells, whereas the increase in amplitude suggested an additional postsynaptic contribution in AII amacrine cells^43,44^. The β-adrenergic receptor agonist isoproterenol mimicked these effects, and the β_1_AR antagonist metoprolol blocked them, supporting a β_1_AR-dependent mechanism (Extended Data Fig. 4).

To test evoked transmission, we optogenetically activated ChR2-expressing rod bipolar cells while recording excitatory postsynaptic currents in AII amacrine cells (Fig. 2j), using a recently established approach that enables stable, long-term analysis of neuromodulation at the rod bipolar cell→AII amacrine cell synapse^45^. Rod bipolar cell stimulation evoked a fast component, reflecting synchronous release, followed by a slower component associated with asynchronous or delayed release (Fig. 2j). Norepinephrine increased the fast component, shortened the time to peak, and suppressed the slow component (Figs. 2j-m). Isoproterenol produced a similar effect, whereas metoprolol blocked the enhancement of the fast component induced by either isoproterenol or norepinephrine (Extended Data Fig. 5).

A previous study suggested that α_2_-adrenergic receptors might modulate rat rod bipolar cell function^46^. However, consistent with our single-cell transcriptomic analysis (Fig. 1c), RNAscope experiments showed that *Adra2a*, *Adra2b*, and *Adra2c* mRNAs were barely detectable in rat or mouse rod bipolar cells (Extended Data Fig. 6a). Moreover, the α_2_ agonist brimonidine had no effect on evoked excitatory postsynaptic currents recorded from mouse AII amacrine cells, whereas RNAscope confirmed *Adrb1* expression in rat rod bipolar cells (Extended Data Figs. 6b,c).

Together, these results show that β_1_AR activation does not simply increase all forms of transmitter release. Instead, it shifts rod bipolar cell output towards faster and more temporally precise transmission. This operation is well suited to the scotopic pathway, where sensitivity must be enhanced without amplifying delayed synaptic noise.

### G_s_-cAMP-PKA pathway mediates the effect

The increase in miniature event amplitude suggested that β_1_AR might also act postsynaptically in AII amacrine cells. Analysis of mouse amacrine cell transcriptomic data (GEO accession number: GSE149715)^47^ revealed *Adrb1* expression in AII amacrine cells but not in starburst amacrine cells (Fig. 3a). RNAscope confirmed *Adrb1* expression in glycinergic amacrine cells positioned near the inner plexiform layer, consistent with AII identity (Fig. 3b). Triple labelling in rat retina further supported *Adrb1* expression in AII amacrine cells (Extended Data Fig. 6d). Comparative analysis indicated that AII expression of *ADRB1* may be rodent-biased rather than broadly conserved across Euarchontoglires (Extended Data Fig. 6e).

**Fig. 3.**
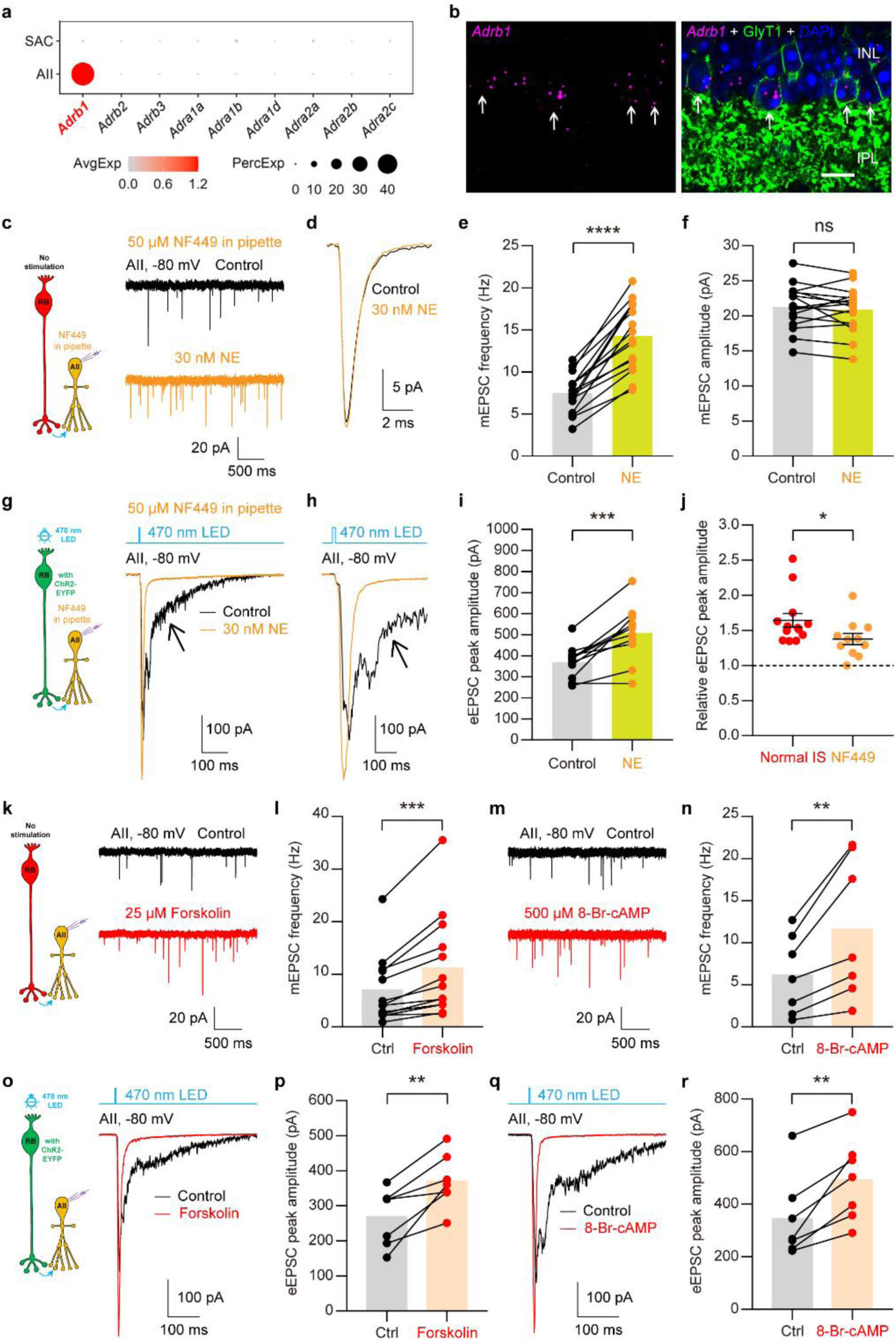
β_1_AR signals through the G_s_-AC-cAMP-PKA pathway. **a**, Single-cell transcriptomic analysis showing *Adrb1* expression in mouse AII amacrine cells but not in starburst amacrine cells (SACs). **b**, RNAscope validation of *Adrb1* expression in GlyT1-positive glycinergic amacrine cells positioned near the IPL (indicated by arrows). DAPI staining is used to label retinal layers. INL, inner nuclear layer; IPL, inner plexiform layer. Scale bar, 10 μm. Images are representative of n = 3 experiments. **c**, Recording configuration for miniature excitatory postsynaptic currents (mEPSCs) from AII amacrine cells with the G_s_ inhibitor NF449 in the patch pipette (left), and representative mEPSCs before and after NE application (right). **d**, Average mEPSC waveforms from the same AII in **c**, showing that NE no longer enhances mEPSC amplitude. **e,f**, Quantification showing that postsynaptic G_s_ inhibition prevents NE-induced enhancement of mEPSC amplitude (**f**; P = 0.4071) but not frequency (**e**; P < 0.0001). n = 16 cells from 6 mice. **g**, Recording configuration for optogenetically evoked rod bipolar cell input to AII amacrine cells with NF449 in the patch pipette (left), and representative evoked excitatory postsynaptic currents (eEPSCs) before and after NE application (right). **h**, Enlarged view of average eEPSCs from the same AII in **g**. **i**, Quantification of the NE effect on eEPSC amplitude during postsynaptic G_s_ inhibition. P = 0.0003. n = 11 cells from 5 mice. **j**, Quantification showing that postsynaptic G_s_ inhibition partially reduces NE-induced enhancement of evoked AII responses. IS: intracellular solution. P = 0.0352. Data are mean ± s.e.m. **k-n**, Activation of adenylyl cyclase with forskolin or PKA with 8-Br-cAMP mimics NE-induced enhancement of mEPSC frequency (P = 0.0002, n = 13 cells from 5 mice for forskolin; P = 0.0095, n = 7 cells from 4 mice for 8-Br-cAMP). **o-r**, Adenylyl cyclase or PKA activation reproduces the NE-induced shift from slow to fast evoked transmission (P = 0.0050 for forskolin; P = 0.0034 for 8-Br-cAMP). n = 7 cells each from 4 and 5 mice, respectively. Paired *t*-test or Mann-Whitney test. *P < 0.05; **P < 0.01; ***P < 0.001; ****P < 0.0001; ns, not significant. Except in **j**, data are presented as individual points with means; error bars are not shown. See also Extended Data Figs. 6-8.

To separate pre- and postsynaptic effects, we included the G_s_ inhibitor NF449 in the patch pipette during AII recordings (Fig. 3c)^48^. Intracellular G_s_ inhibition abolished the norepinephrine-induced increase in miniature event amplitude but did not prevent the increase in miniature event frequency (Figs. 3c-f). Similarly, during evoked recordings, postsynaptic G_s_ inhibition reduced the magnitude of norepinephrine-induced enhancement but preserved the presynaptic shift towards faster rod bipolar cell output (Figs. 3g-j). Thus, β_1_AR acts on both sides of the rod bipolar cell→AII synapse: presynaptically to regulate release and postsynaptically, at least in rodents, to increase AII responsiveness.

Pharmacological manipulation of the canonical β_1_AR pathway supported this model (Extended Data Fig. 7). Activation of adenylyl cyclase or PKA mimicked norepinephrine by increasing miniature event frequency (Figs. 3k-n) and by enhancing the fast component of evoked transmission while suppressing the slow component (Figs. 3o-r and Extended Data Figs. 7b,c). Conversely, inhibition of adenylyl cyclase or PKA prevented the norepinephrine effect (Extended Data Figs. 7d,e). Recordings from rod bipolar cells further showed that norepinephrine depolarized rod bipolar cells without altering evoked depolarization amplitude or voltage-gated calcium currents (Extended Data Fig. 8). Together, these data indicate that β_1_AR engages G_s_-adenylyl cyclase-cAMP-PKA signaling to tune rod bipolar cell output, most likely by altering terminal excitability rather than calcium-channel conductance.

### β_1_AR boosts retinal output

We next asked whether β_1_AR-dependent modulation of the rod bipolar cell→AII synapse propagates to ganglion cells, the output neurons of the retina. In ON α ganglion cells, optogenetic activation of rod bipolar cells evoked excitatory currents that were strongly reduced by blockade of electrical coupling between AII amacrine cells and ON cone bipolar cells (Extended Data Figs. 9a,b), confirming that these responses were mediated by the canonical rod bipolar cell pathway^2,27^. Norepinephrine increased the amplitude and shortened the time to peak of these evoked excitatory currents (Figs. 4a-c), while modestly accelerating their decay (Extended Data Fig. 9c). In current-clamp recordings, norepinephrine increased light-evoked spike number and shortened first spike latency in ON α ganglion cells (Figs. 4d-g), while modestly depolarizing their resting membrane potential (Extended Data Fig. 9d).

**Fig. 4.**
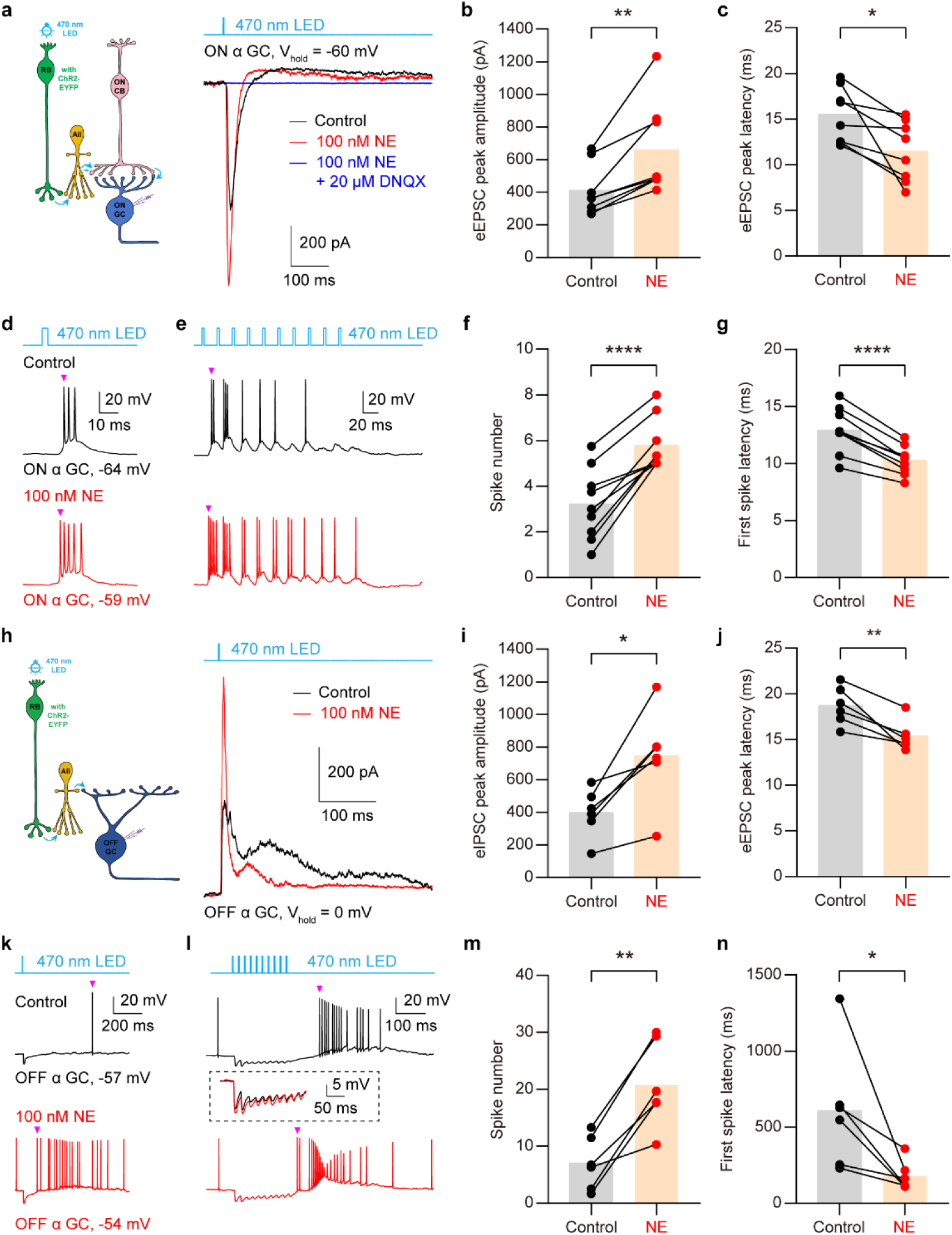
β_1_AR activation enhances retinal output. **a**, Recording configuration for evoked excitatory postsynaptic currents (eEPSCs) from ON α ganglion cells during optogenetic stimulation of rod bipolar cells (left), and representative ON α ganglion cell eEPSCs before and after norepinephrine (NE) application and subsequent blockade by DNQX (right). **b,c**, Quantification of ON α ganglion cell eEPSC peak amplitude (P = 0.0078) and peak latency (P = 0.0154). n = 8 cells from 5 mice. **d-g**, Current-clamp recordings showing that NE increases light-evoked spiking (P < 0.0001) and shortens first spike latency (P < 0.0001) in ON α ganglion cells. Arrowheads indicate the first spike following light stimulation. n = 9 cells from 5 mice. **h**, Recording configuration for inhibitory postsynaptic currents (eIPSCs) from OFF α ganglion cells during optogenetic stimulation of rod bipolar cells (left), and representative OFF α ganglion cell eIPSCs before and after NE application (right). **i,j**, Quantification of OFF α ganglion cell eIPSC peak amplitude (P = 0.0108) and peak latency (P = 0.0062). n = 6 cells from 5 mice. **k-n**, Current-clamp recordings showing that NE enhances light-evoked hyperpolarization (inset) and post-light spiking (P = 0.0016) and shortens first spike latency (P = 0.0413) in OFF α ganglion cells. Arrowhead indicates the first spike following light stimulation. n = 6 cells from 6 mice. Paired *t*-test or Wilcoxon test. *P < 0.05; **P < 0.01; ****P < 0.0001. Data are presented as individual points with means; error bars are not shown. See also Extended Data Fig. 9.

OFF α ganglion cells receive scotopic input through glycinergic inhibition from AII amacrine cells^2,27^. Optogenetically evoked inhibitory postsynaptic currents in OFF α ganglion cells were abolished by blocking glutamatergic transmission from rod bipolar cells to AII amacrine cells (Extended Data Figs. 9e,f), confirming their dependence on the rod bipolar cell pathway. Consistent with enhanced rod bipolar cell→AII signaling, norepinephrine increased the peak amplitude and shortened the time to peak of evoked inhibitory currents in OFF α ganglion cells (Figs. 4h-j), while modestly accelerating their decay (Extended Data Fig. 9g). In current clamp, norepinephrine enhanced light-evoked hyperpolarization, increased post-light spiking and shortened first spike latency (Figs. 4k-n), without altering the resting membrane potential (Extended Data Fig. 9h). Together, these results show that β_1_AR activation strengthens and accelerates both ON and OFF retinal output downstream of the rod bipolar cell pathway.

Thus, the synaptic modulation identified at the rod bipolar cell terminal is propagated through the inner retina to alter ganglion cell output. β_1_AR activation enhances both the gain and timing of scotopic retinal signaling, providing a circuit-level mechanism by which adrenergic state can improve low-light visual responses.

### *Adrb1* deletion abolishes night-vision gain

To test the in vivo consequence of β_1_AR activation, we measured scotopic electroretinograms after intravitreal injection of norepinephrine or vehicle into opposite eyes of the same mouse (Fig. 5a). Norepinephrine increased dark-adapted b-wave amplitudes across a broad range of flash intensities (Figs. 5a,b). Isoproterenol produced a similar enhancement, consistent with β-adrenergic modulation of retinal responses (Fig. 5c).

**Fig. 5.**
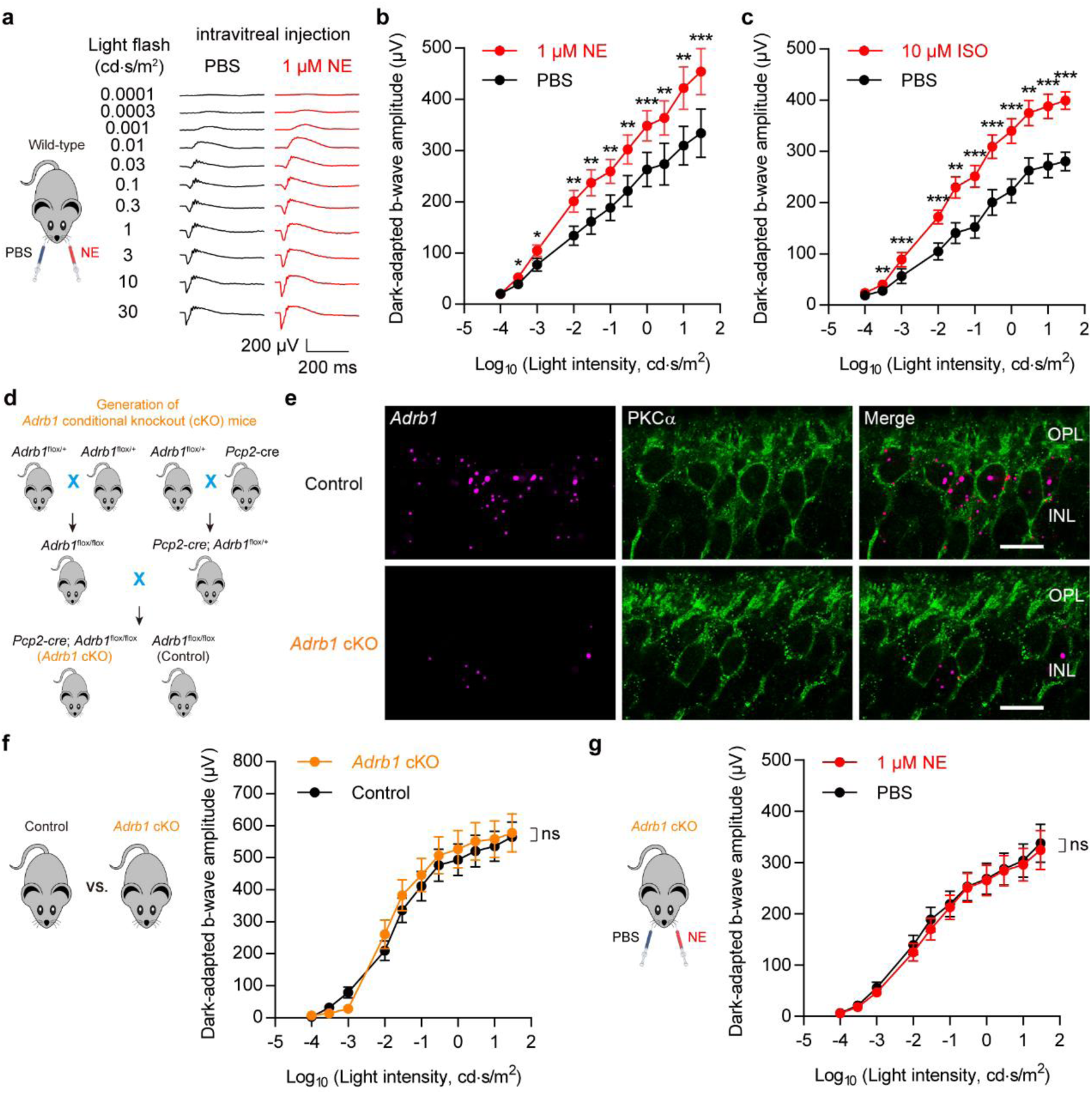
Rod bipolar cell *Adrb1* is required for adrenergic enhancement of scotopic vision. **a**, Experimental design for scotopic electroretinography (ERG) after intravitreal injection of norepinephrine (NE) or PBS into opposite eyes of the same mouse (left), and average scotopic ERG traces from eight wild-type mice (right). **b,c**, Intravitreal injection of NE (**b**) or isoproterenol (ISO; **c**) increases dark-adapted b-wave amplitudes across flash intensities. n = 8 mice for each group. **d**, Strategy for generating rod-bipolar-cell-specific *Adrb1* conditional knockout (cKO) mice. **e**, RNAscope validation showing loss of *Adrb1* transcripts in cKO rod bipolar cells. OPL, outer plexiform layer; INL, inner nuclear layer. Scale bar, 10 μm. Images are representative of n = 3 experiments. **f**, Baseline scotopic ERG responses are unchanged in *Adrb1* cKO mice. N = 12 mice for each group. **g**, Intravitreal NE fails to enhance dark-adapted b-wave amplitudes in *Adrb1* cKO mice. n = 16 mice. Paired or unpaired *t*-tests. *P < 0.05; **P < 0.01; ***P < 0.001; ns, not significant. Data are mean ± s.e.m. See also Extended Data Fig. 10.

We then generated mice in which *Adrb1* was conditionally deleted from rod bipolar cells (Fig. 5d). RNAscope confirmed loss of *Adrb1* transcripts from rod bipolar cells in conditional knockout retinas (Fig. 5e). Baseline scotopic electroretinograms were indistinguishable between conditional knockout mice and control littermates (Fig. 5f), and pharmacological β_1_AR blockade did not significantly affect baseline responses in wild-type mice (Extended Data Fig. 10). Thus, β_1_AR is not required for basal rod bipolar cell pathway function.

The critical difference emerged under adrenergic stimulation. In control mice, norepinephrine enhanced scotopic b-wave amplitudes, whereas in rod-bipolar-cell-specific *Adrb1* conditional knockout mice this enhancement was abolished (Fig. 5g). These results establish a causal requirement for rod bipolar cell β_1_AR in norepinephrine-dependent enhancement of scotopic retinal function. β_1_AR therefore acts not as a constitutive component of baseline vision, but as an inducible gain-control mechanism that can be recruited when norepinephrine levels rise, such as during arousal or stress.

### *ADRB1* is redeployed in tree shrew cones

The conservation of *ADRB1* expression in rod bipolar cells across sampled Euarchontoglires suggested that this module may have been maintained because it improves vision under low-light conditions. We therefore examined the tree shrew, a Euarchontoglires species with a highly cone-dominated retina (Fig. 6a) and a diurnal visual ecology^37^.

**Fig. 6.**
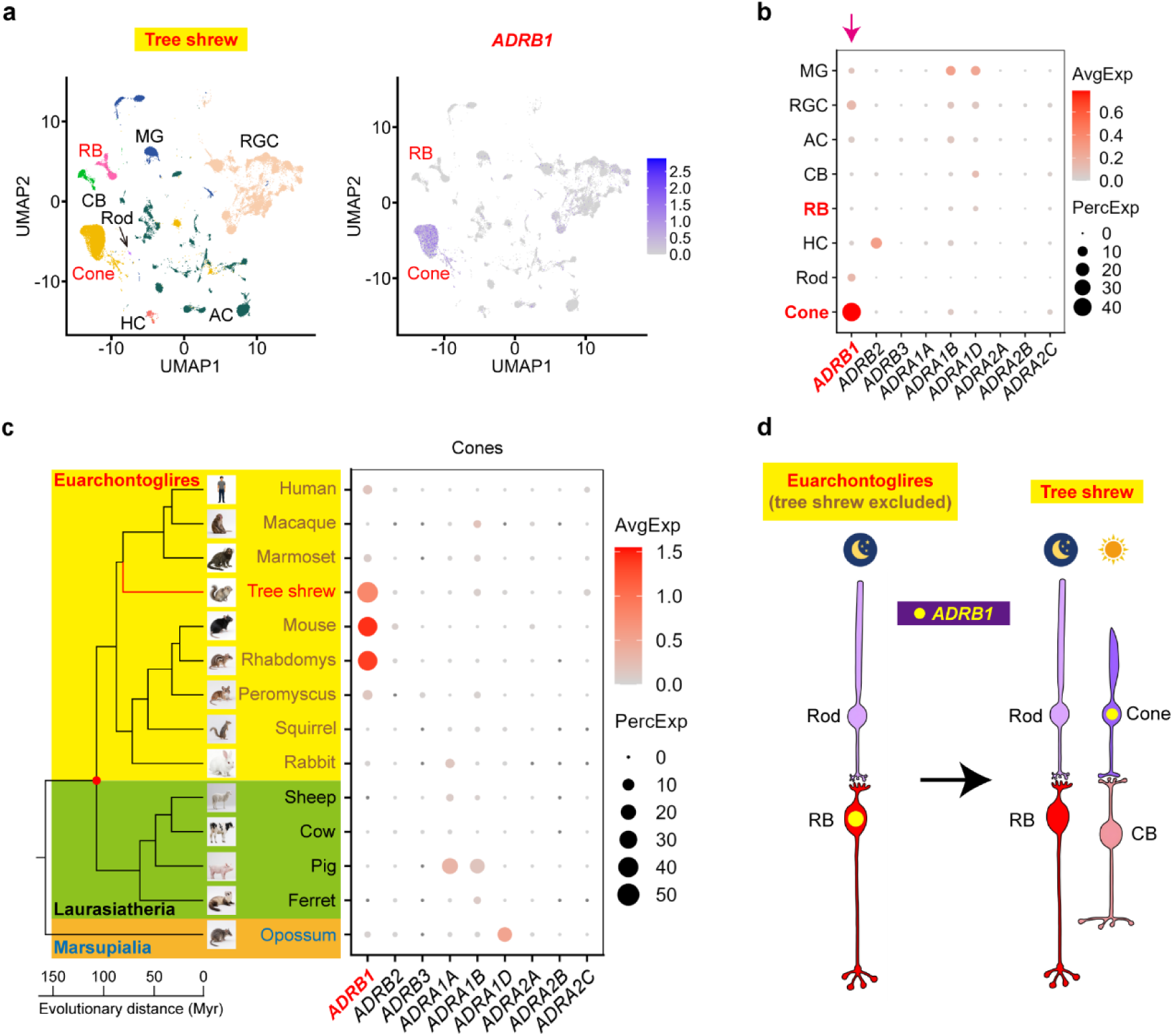
*ADRB1* is redeployed to cones in the diurnal tree shrew. **a**, UMAP visualization of tree shrew retinal cells, with *ADRB1* expression shown on the right. Note that the number of cones (n = 13157 cells) far exceeds that of rods (n = 166 cells; arrow) in the tree shrew single-cell RNA sequencing data. **b**, Dot plot of *ADRB1* expression (arrow) across tree shrew retinal cell types. **c**, Cross-species comparison of cone *ADRB1* expression showing enrichment in tree shrew and selected Euarchontoglires, but not in sampled Laurasiatheria or Marsupialia. **d**, Model for evolutionary redeployment of an adrenergic module. In most sampled Euarchontoglires, β_1_AR, encoded by *ADRB1*, is positioned in rod bipolar cells to enhance scotopic gain; in the cone-dominated tree shrew retina, the module is shifted to cones, where it may tune photopic or cone-mediated vision.

Unlike other sampled Euarchontoglires, the tree shrew showed little or no *ADRB1* expression in rod bipolar cells (Figs. 6a,b). Instead, *ADRB1* was strongly expressed in cone photoreceptors (Figs. 6a-c). This pattern was not observed in sampled Laurasiatheria, whose cones lacked comparable *ADRB1* expression (Fig. 6c). The tree shrew therefore does not simply lack the adrenergic module; rather, it appears to have reassigned this module from the rod bipolar cell pathway to the photoreceptor class that dominates its visual ecology.

This finding reframes *ADRB1* as an evolutionarily mobile neuromodulatory module rather than a fixed marker of a single cell type. In most sampled Euarchontoglires, β_1_AR is positioned in rod bipolar cells, where it enhances low-light retinal gain. In the diurnal, cone-dominated tree shrew, the same receptor is redeployed to cones, where it may instead modulate daylight or cone-mediated vision (Fig. 6d). Direct physiological studies in tree shrew will be required to test this functional prediction, but the expression shift provides a natural example of cell-type reassignment of a conserved neuromodulatory mechanism.

## Discussion

We identify a retinal adrenergic module associated with one of the deepest splits in placental mammalian evolution. *ADRB1* is expressed in rod bipolar cells across sampled Euarchontoglires but is absent from homologous cells in sampled Laurasiatheria and Marsupialia retina (Fig. 1 and Extended Data Figs. 1-3). In mice, this molecular difference has a clear functional consequence: β_1_AR activation enhances rod bipolar cell synaptic output, strengthens downstream retinal signaling, and increases scotopic responses in vivo (Figs. 2-5 and Extended Data Figs. 4-10). Conditional deletion of *Adrb1* from rod bipolar cells abolishes norepinephrine-induced enhancement without impairing baseline retinal function (Fig. 5 and Extended Data Fig. 10). These results connect a cell-type-specific evolutionary transcriptomic signature to synaptic mechanism and visual output.

A central implication of this work is that sensory evolution can act through neuromodulatory access to conserved neural circuits. The rod bipolar cell pathway is an ancient and highly conserved circuit for dim-light vision^2,3,14,21^. Rather than altering the core wiring of this pathway, Euarchontoglires appear to have coupled it to adrenergic gain control by expressing β_1_AR at a key synaptic node. Such a mechanism is economical: it preserves baseline retinal computation while allowing behavioral state to enhance sensitivity and temporal precision when needed.

The location of β_1_AR in rod bipolar cells may be functionally important. Rod photoreceptors operate near the physical limit of photon detection, and indiscriminate amplification at the photoreceptor level could amplify intrinsic noise^25,28,49^. By placing gain control at the rod bipolar cell terminal, the retina can enhance downstream signaling after initial photon detection while improving the timing of synaptic output^25,28–30,49^. Our recordings support this idea: β_1_AR activation enhances synchronous release while suppressing asynchronous/delayed release, a combination that should increase signal gain and reduce temporally dispersed noise in the scotopic pathway.

The evolutionary interpretation should be considered in light of species sampling. The available Laurasiatheria data include representatives of Cetartiodactyla (pig, cow, yellow cattle, sheep) and Carnivora (ferret, cat), but not Chiroptera, Perissodactyla, Eulipotyphla, or Pholidota^5,7–9^. Additional single-cell datasets from these lineages will be necessary to determine whether absence of rod bipolar cell *ADRB1* is universal across Laurasiatheria. The opossum outgroup shows little or no rod bipolar cell *ADRB1* expression, tentatively favoring a gain of rod bipolar cell *ADRB1* in the Euarchontoglires lineage rather than loss in Laurasiatheria. However, broader sampling of Marsupialia, Afrotheria, and Xenarthra will be required to resolve the ancestral state^7–9^.

The tree shrew provides an important qualification and conceptual extension. As a Euarchontoglires species with a cone-dominated retina^37^, tree shrew has shifted *ADRB1* expression from rod bipolar cells to cones (Fig. 6). This result argues against a simple “Euarchontoglires equals rod bipolar cell *ADRB1*” rule. Instead, it suggests that the adrenergic module is evolutionarily redeployable. In visual ecologies where rod bipolar cell signaling is central, the module is positioned in rod bipolar cells; in a cone-dominated diurnal lineage, it is repositioned in cones. This cell-type reassignment may represent a general strategy by which evolution tunes conserved neural circuits to ecological demand.

Our findings also raise mechanistic questions. The regulatory elements that drive rod bipolar cell *ADRB1* expression in most Euarchontoglires, and cone *ADRB1* expression in tree shrew, remain unknown. Comparative epigenomic and cis-regulatory analyses could identify enhancers responsible for this cell-type specificity and test whether they are absent, inactive, or repurposed in Laurasiatheria^50^. Similarly, physiological studies in tree shrew are needed to determine whether cone β_1_AR enhances cone synaptic output, photopic sensitivity, or state-dependent daylight vision.

More broadly, this study illustrates how cross-species single-cell transcriptomics can move beyond cataloguing cell types to reveal functional principles of sensory evolution. By integrating comparative transcriptomics with neural circuit physiology and conditional genetics, we show that a single neuromodulatory receptor can provide an evolutionary link between molecular identity, synaptic computation, and visual ecology. Such mobile neuromodulatory modules may be common across sensory systems, offering a flexible route by which evolution adapts conserved neural circuits to the demands of different environments.

## Extended Data figures and figure legends

**Extended Data Fig. 1.**
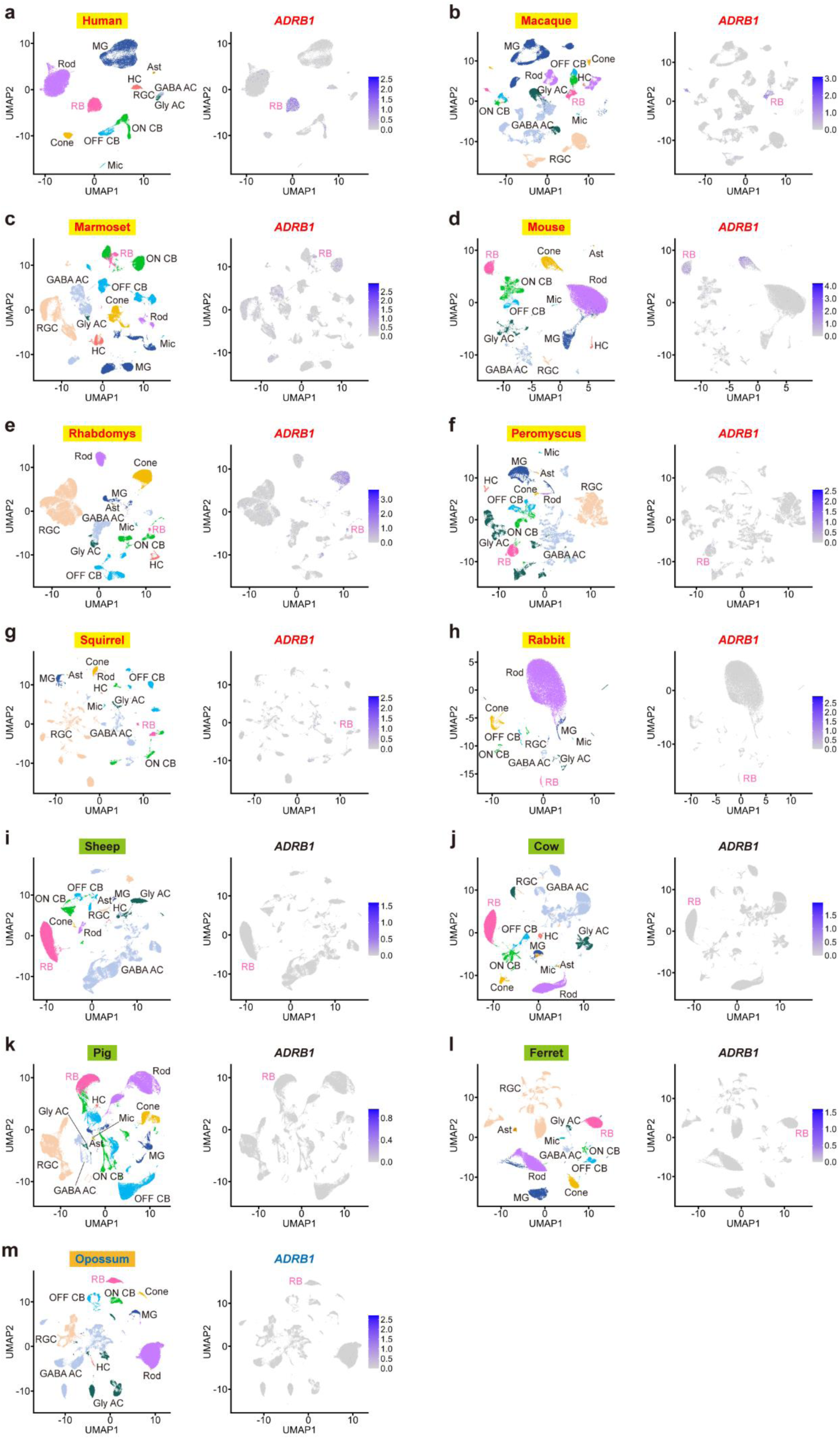
UMAP visualization of *ADRB1* expression across species. **a-m**, Uniform manifold approximation and projection (UMAP) embeddings of single-cell RNA-sequencing data from Euarchontoglires, Laurasiatheria, and Marsupialia retinas. Cell numbers (per species): human (32198), macaque (69503), marmoset (55981), mouse (40970), rhabdomys (63452), peromyscus (46499), squirrel (22840), rabbit (27048), sheep (67767), cow (78283), pig (64999), ferret (47720), and opossum (78526). Right panels show *ADRB1* expression levels. See Fig. 1a for cell-type annotations. Related to Fig. 1.

**Extended Data Fig. 2.**
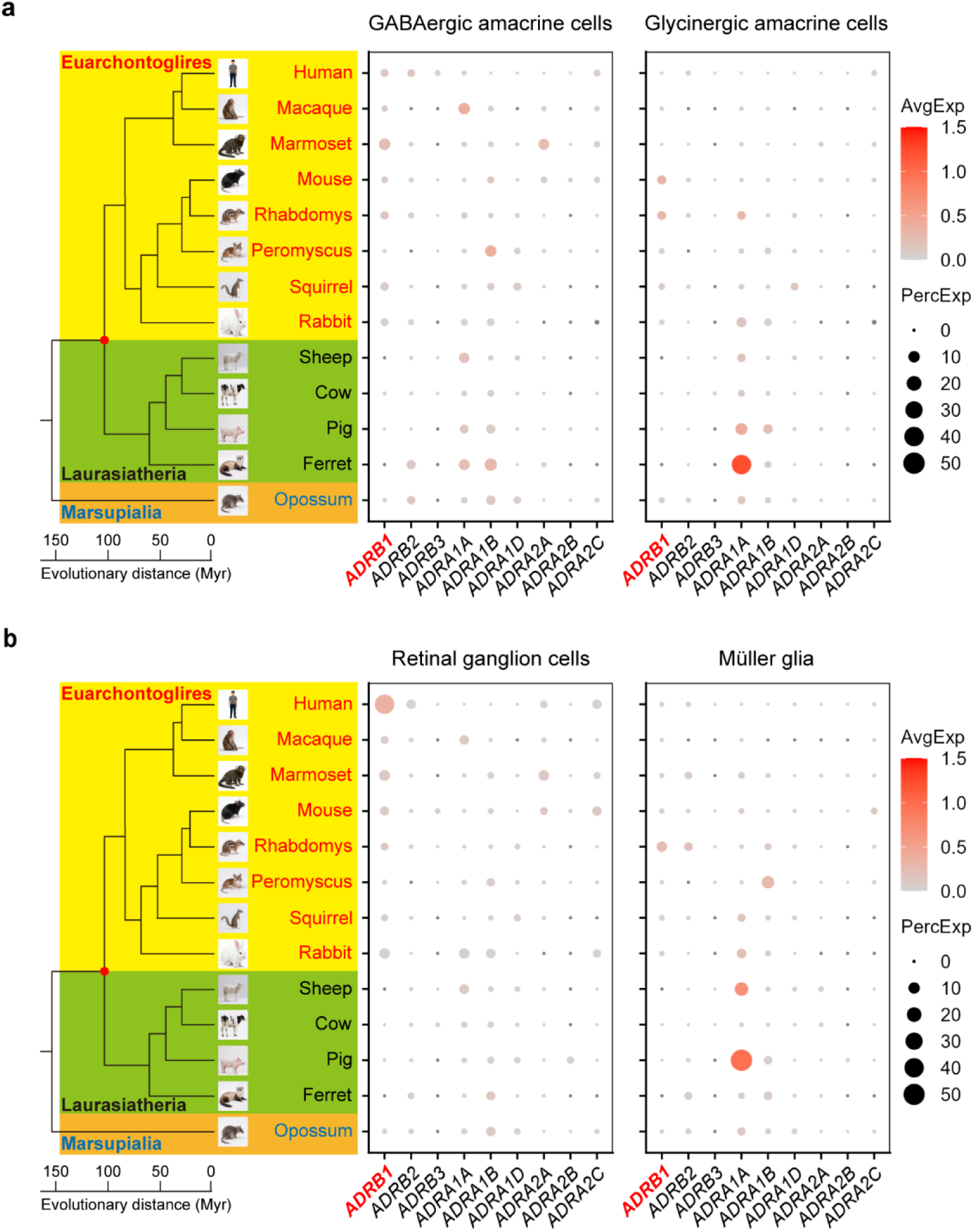
Adrenergic receptor expression in amacrine cells, ganglion cells, and Müller glia. **a**, Dot plots showing expression of adrenergic receptor genes in GABAergic and glycinergic amacrine cells across 13 species. **b**, Dot plots showing expression of adrenergic receptor genes in retinal ganglion cells and Müller glia. Phylogenetic relationships and divergence times are shown on the left. Dot size indicates the fraction of expressing cells and color indicates average normalized expression. No cladistic pattern comparable to *ADRB1* expression in rod bipolar cells is observed. Myr, million years. Related to Fig. 1.

**Extended Data Fig. 3.**
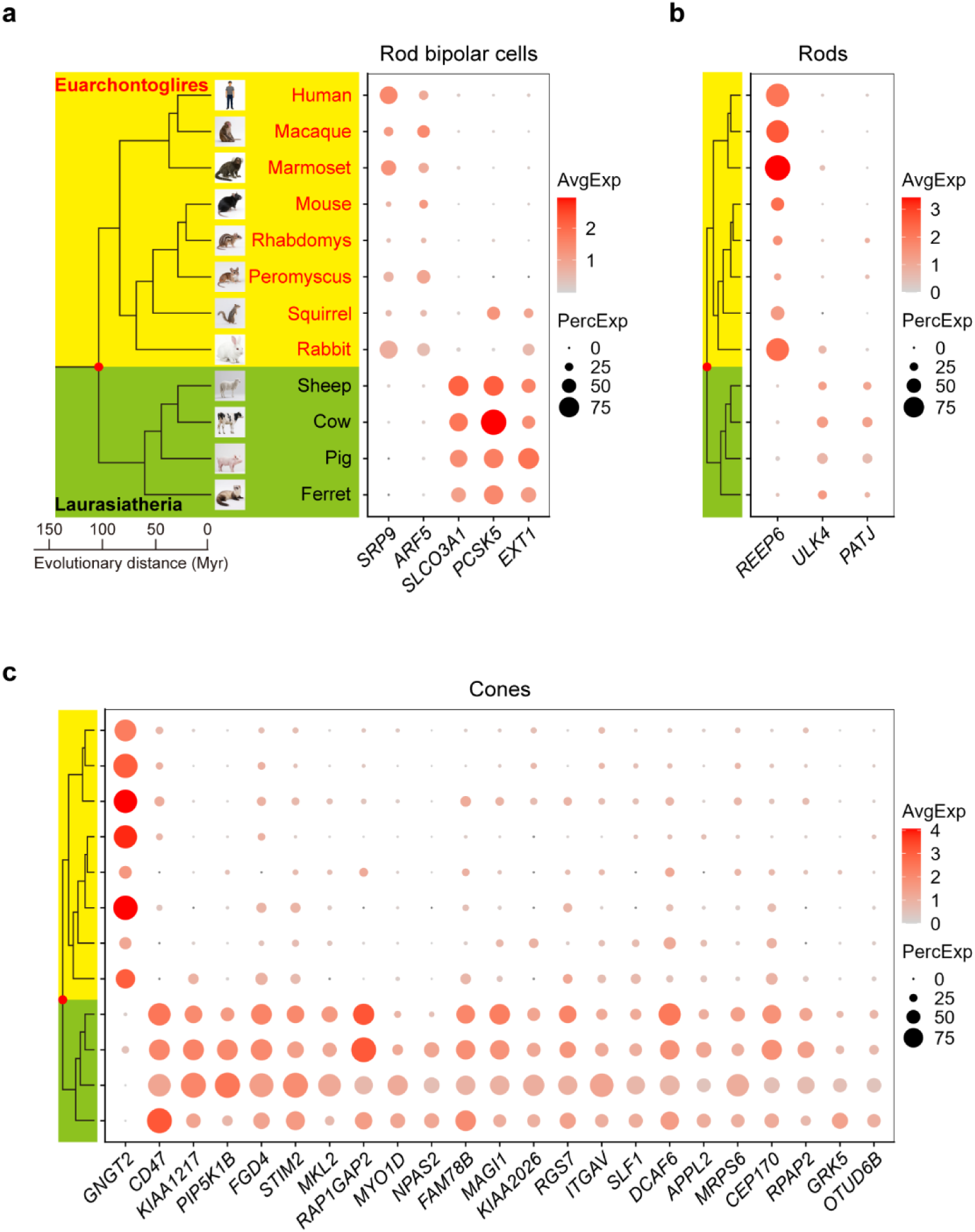
Additional superorder-associated retinal genes. **a**, Dot plots showing genes preferentially expressed in rod bipolar cells of Euarchontoglires or Laurasiatheria, including *SRP9* and *SLCO3A1*. **b**, Genes enriched in rods of Euarchontoglires or Laurasiatheria, including *REEP6* and *ULK4*. **c**, Genes enriched in cones of Euarchontoglires or Laurasiatheria, including *GNGT2* and *CD47*. Species are ordered as in **a**. See Fig. 1c for the *ADRB1* comparison. Related to Fig. 1.

**Extended Data Fig. 4.**
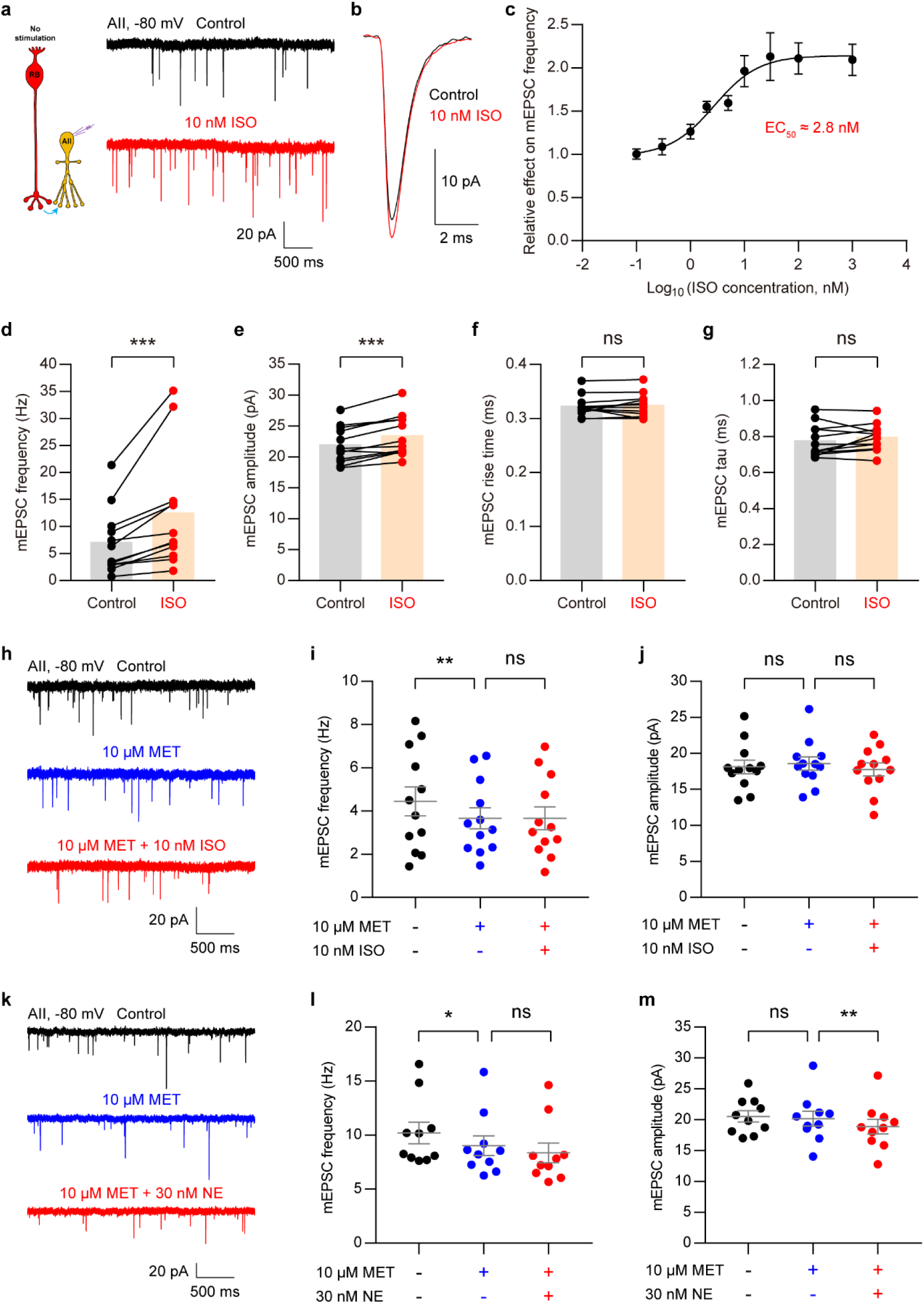
Pharmacological validation of β_1_AR-dependent modulation of spontaneous release. **a**, Recording configuration for miniature excitatory postsynaptic currents (mEPSCs) from an AII amacrine cell (left), and representative mEPSCs under control and isoproterenol (ISO; β-adrenergic receptor agonist; 10 nM) conditions (right). **b**, Average mEPSC waveforms from the same AII in **a** under control and ISO conditions. **c**, Dose-response relationship for the effect of ISO on mEPSC frequency. EC_50_ ≈ 2.8 nM. **d-g**, Quantification of ISO effects on mEPSC frequency (P = 0.0005), amplitude (P = 0.0002), rise time (P = 0.4238), and decay tau (P = 0.2334). n = 12 cells from 6 mice. Paired *t*-test. Data are presented as individual points with means; error bars are not shown. **h**, Representative mEPSCs showing that metoprolol (MET; β_1_AR antagonist; 10 μM) blocks ISO effects. **i,j**, Quantification of MET and ISO effects on mEPSC frequency (**i**; control vs. MET, P = 0.0084; MET vs. MET + ISO, P > 0.9999) and amplitude (**j**; control vs. MET, P = 0.6322; MET vs. MET + ISO, P = 0.2647). n = 12 cells from 5 mice. RM-ANOVA, Sidak’s multiple-comparisons test. **k**, Representative mEPSCs showing that MET blocks NE effects. **l,m**, Quantification of MET and NE effects on mEPSC frequency (**l**; control vs. MET, P = 0.0146; MET vs. MET + NE, P > 0.9999; Friedman test, Dunn’s multiple-comparisons test) and amplitude (**m**; control vs. MET, P = 0.8282; MET vs. MET + NE, P = 0.0042; RM-ANOVA, Sidak’s multiple-comparisons test). n = 10 cells from 4 mice. *P < 0.05; **P < 0.01; ***P < 0.001; ns, not significant. Unless otherwise indicated, data are presented as mean ± s.e.m. Related to Fig. 2.

**Extended Data Fig. 5.**
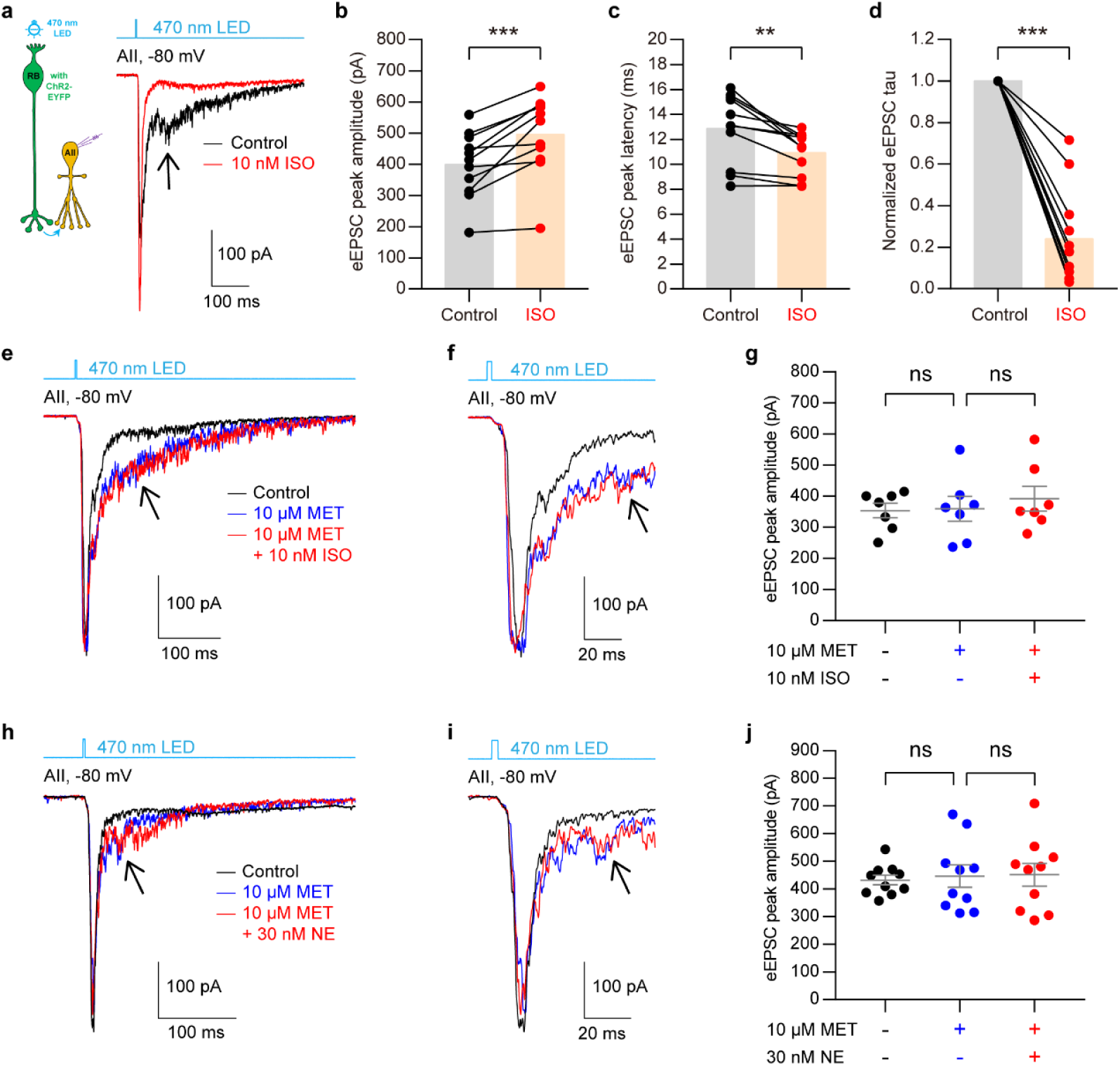
Pharmacological validation of β_1_AR-dependent modulation of evoked release. **a**, Recording configuration for evoked excitatory postsynaptic currents (eEPSCs) from AII amacrine cells during optogenetic stimulation of rod bipolar cells (left), and representative eEPSCs showing fast and slow components under control and isoproterenol (ISO; β-adrenergic receptor agonist; 10 nM) conditions (right). **b**-**d**, Quantification of ISO effects on eEPSC peak amplitude (P = 0.0007), peak latency (P = 0.0023), and decay tau (P = 0.0010). n = 11 cells from 6 mice. Paired *t*-test or Wilcoxon test. Data are presented as individual points with means; error bars are not shown. **e**, MET blocks ISO effects. **f**, Enlarged view of average eEPSCs from the same AII in **e**. **g**, Quantification of MET and ISO effects on eEPSC peak amplitude (control vs. MET, P = 0.9663; MET vs. MET + ISO, P = 0.4142). n = 7 cells from 3 mice. RM-ANOVA, Sidak’s multiple-comparisons test. **h**, Representative eEPSCs showing that MET enhances the slow component and blocks the NE effect (right). **i**, Enlarged view of average eEPSCs from the same AII in **h**. **j**, Quantification of MET and NE effects on eEPSC peak amplitude (control vs. MET, P = 0.8563; MET vs. MET + NE, P = 0.9772). n = 10 cells from 5 mice. RM-ANOVA, Sidak’s multiple-comparisons test. **P < 0.01; ***P < 0.001; ns, not significant. Unless otherwise indicated, data are presented as mean ± s.e.m. Related to Fig. 2.

**Extended Data Fig. 6.**
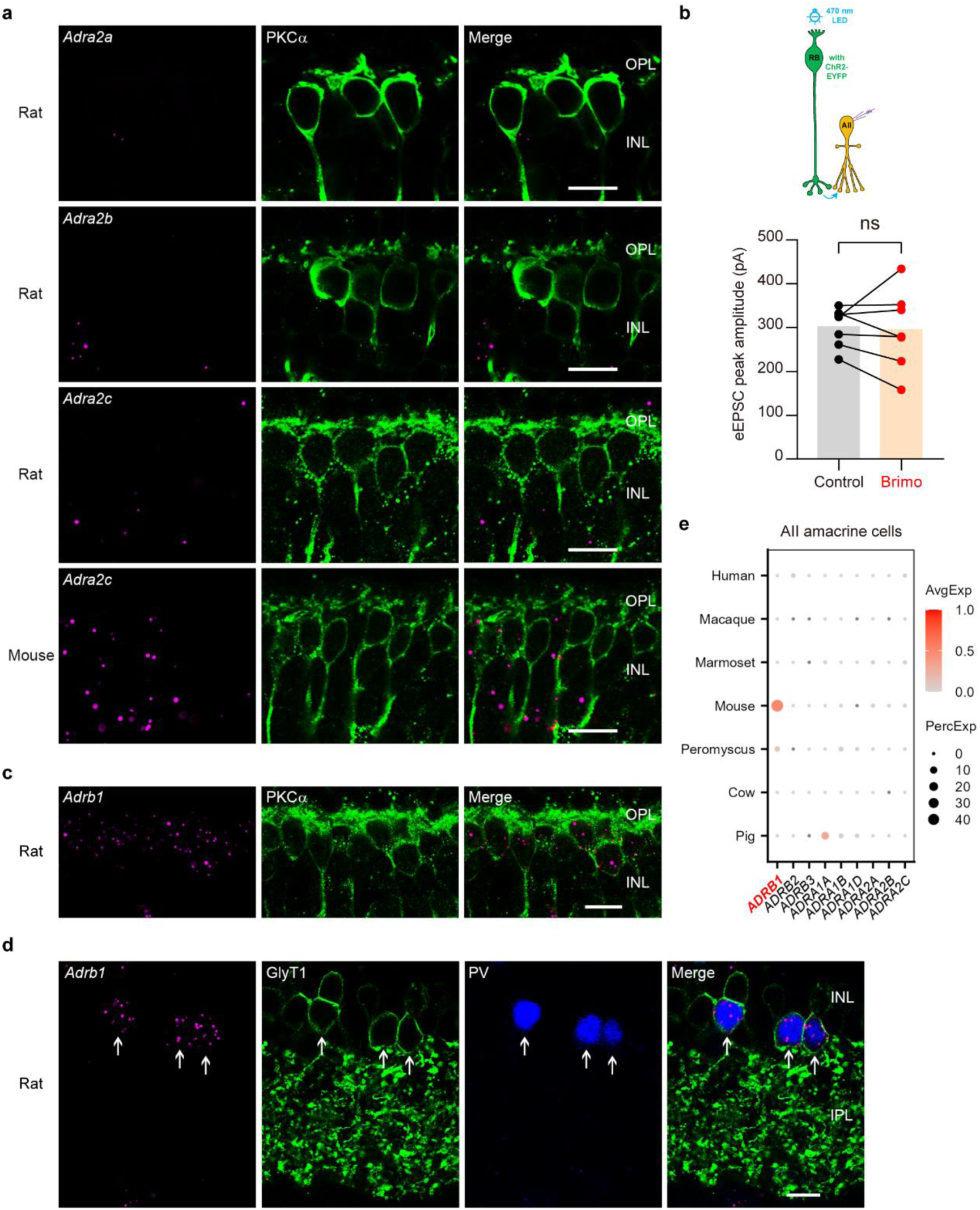
*Adrb1*, but not α_2_-adrenegic receptor mRNAs, are expressed in rodent rod bipolar cells and AII amacrine cells. **a**, RNAscope co-labelling of α_2_-adrenegic receptor mRNAs (*Adra2a*, *Adra2b*, *Adra2c*) with PKCα in rat and mouse retinas. PKCα labels rod bipolar cells. **b**, Recording configuration for evoked excitatory postsynaptic currents (eEPSCs) from AII amacrine cells with optogenetic stimulation of rod bipolar cells (up), and quantification showing that the α_2_-adrenergic receptor agonist brimonidine (Brimo; 10 μM) has no effect on eEPSC peak amplitude. P = 0.7862. n = 7 cells from 3 mice. **c**, RNAscope co-labelling of *Adrb1* and PKCα in rat retina confirms *Adrb1* expression in rod bipolar cells. **d**, Triple RNAscope labelling of *Adrb1*, GlyT1, and parvalbumin (PV) in rat retina. Arrows indicate AII amacrine cells. **e**, Dot plots showing *Adrb1* expression in AII amacrine cell clusters from seven species. *Adrb1* expression is detected in mouse and weakly in Peromyscus. Scale bars, 10 μm. ns, not significant. Paired *t*-test. Data are mean ± s.e.m. Images in **a**, **c**, and **d** are representative of n ≥ 3 experiments. Related to Figs. 2,3.

**Extended Data Fig. 7.**
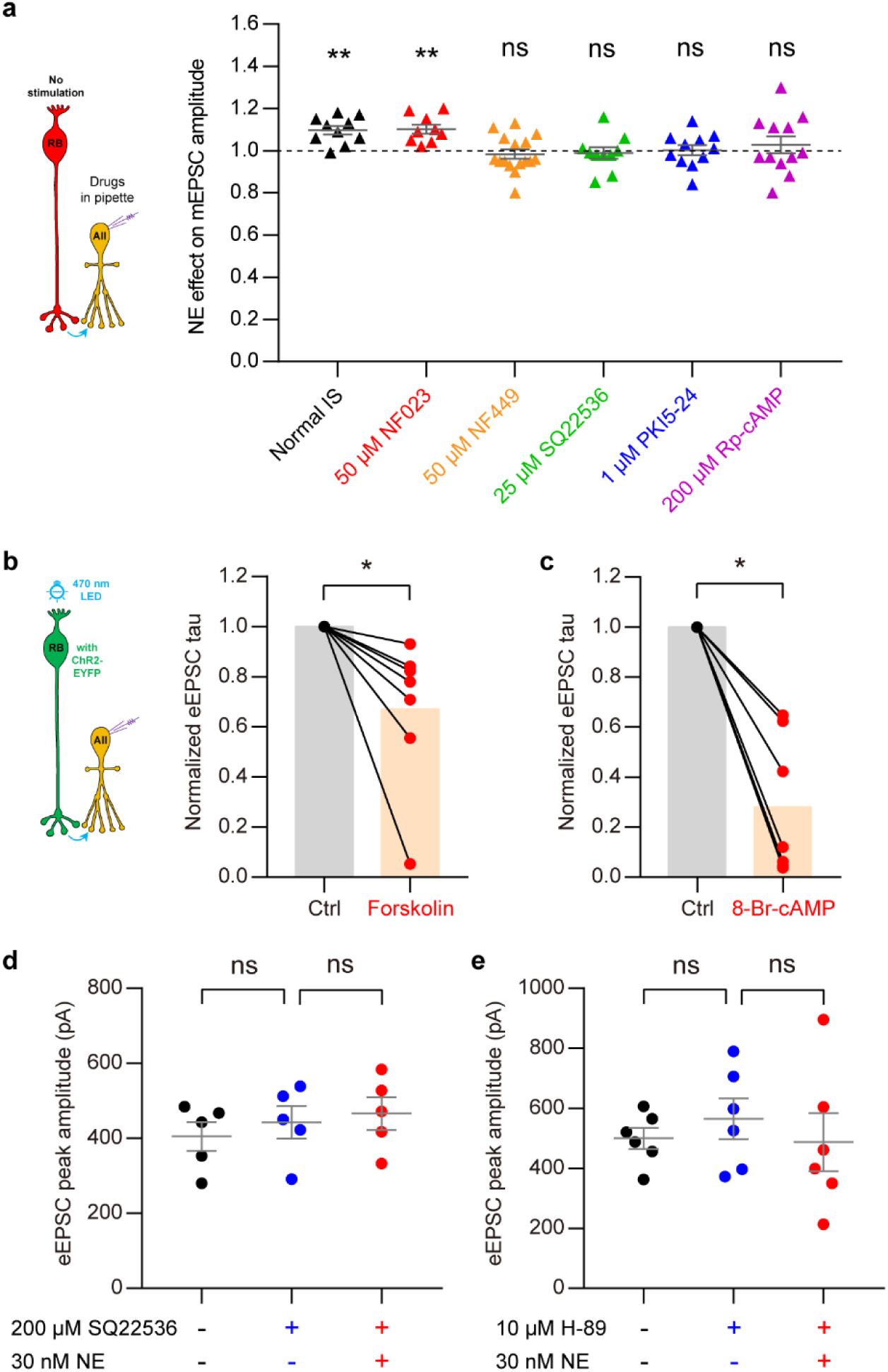
Pre- and postsynaptic β_1_AR signaling through the G_s_-AC-cAMP-PKA pathway. **a**, Recording configuration for miniature excitatory postsynaptic currents (mEPSCs) from AII amacrine cells with different inhibitors in the patch pipette (left). Intracellular inhibition of G_s_ with NF449, adenylyl cyclase with SQ22536 or PKA with PKI5-24 or Rp-cAMP blocks the norepinephrine (NE)-induced increase in mEPSC amplitude, whereas inhibition of G_i_ with NF023 does not (right). Relative NE effects with normal internal solution (IS), NF023, NF449, SQ22536, PKI5-24, and Rp-cAMP: P = 0.0010, 0.0015, 0.4575, 0.6947, 0.9134, and 0.5032; n = 10, 9, 16, 10, 11, and 12 cells. **b,c**, Recording configuration for evoked excitatory postsynaptic currents (eEPSCs) from AII amacrine cells with optogenetic stimulation of rod bipolar cells (**b**, left). Membrane-permeable forskolin (adenylyl cyclase activator; 25 μM; **b**, right) and 8-Br-cAMP (PKA activator; 500 μM; **c**) reduce the decay tau of eEPSCs (P = 0.0156 for forskolin; P = 0.0156 for 8-Br-cAMP). n = 7 cells for each group from 4 and 5 mice, respectively. Wilcoxon test. Data are presented as individual points with means; error bars are not shown. **d,e**, The adenylyl cyclase inhibitor SQ22536 and the PKA inhibitor H-89 prevent NE-induced enhancement of eEPSCs (control vs. SQ22536, P = 0.3690; SQ22536 vs. SQ22536 + NE, P = 0.5072; control vs. H-89, P = 0.4589; H-89 vs. H-89 + NE, P = 0.4248). n = 5 and 6 cells from 3 mice each, respectively. RM-ANOVA, Sidak’s multiple-comparisons test. Data are mean ± s.e.m. *P < 0.05; **P < 0.01; ns, not significant. Related to Fig. 3.

**Extended Data Fig. 8.**
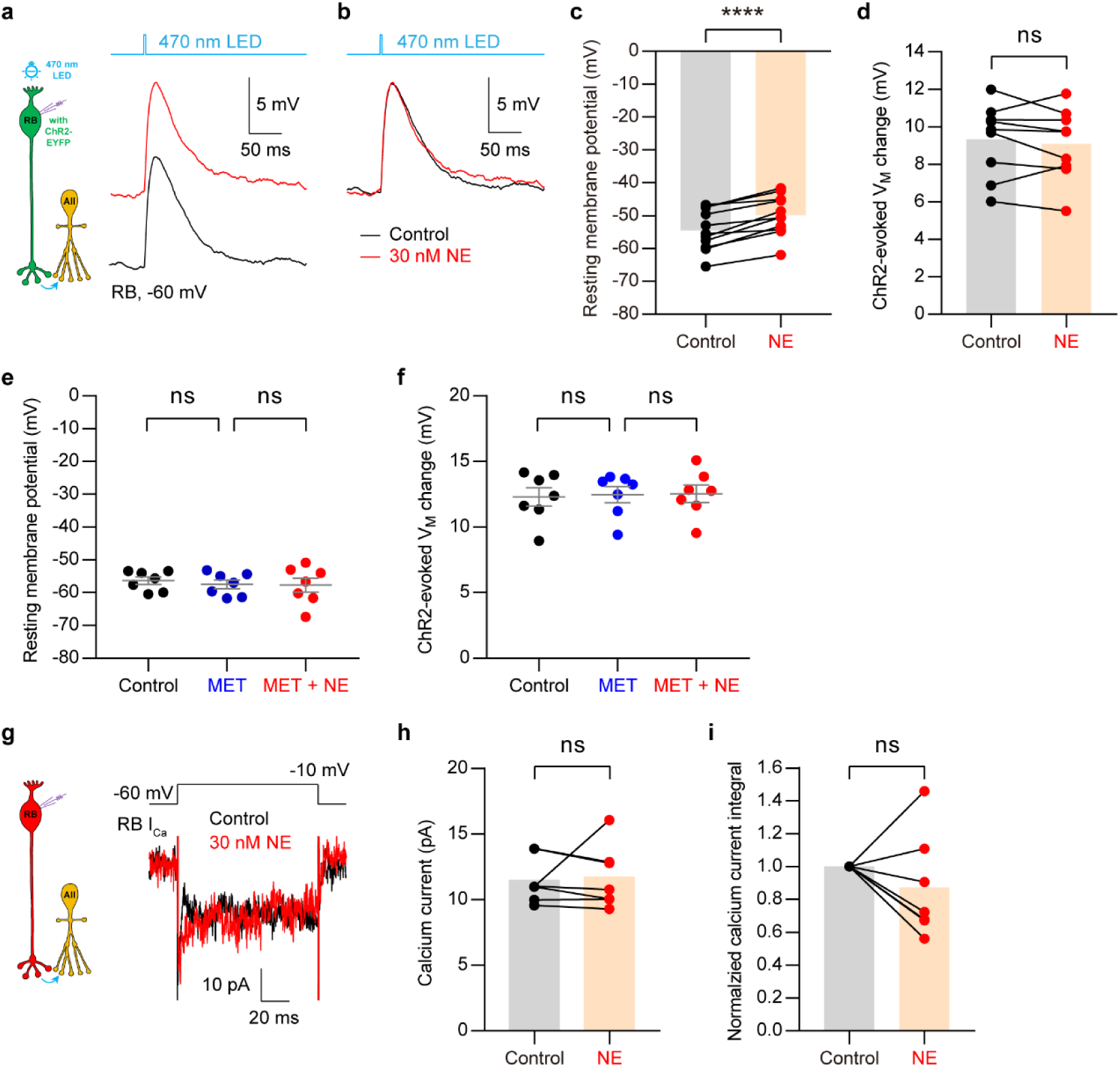
β_1_AR activation depolarizes rod bipolar cells without changing calcium currents. **a**, Recording configuration for rod bipolar (RB) cell membrane potential measurements with optogenetic stimulation (left), and representative traces showing depolarization induced by 30 nM NE (right). **b**, Offset resting membrane potentials from the same RB shown in **a** under control and NE conditions. **c,d**, Quantification of NE effects on resting membrane potential (P < 0.0001; n = 11 cells from 8 mice) and ChR2-evoked depolarization (P = 0.4180; n = 9 cells from 7 mice). **e,f**, MET blocks NE-induced depolarization (control vs. MET, P = 0.2979; MET vs. MET + NE, P = 0.9636) but has no effect on ChR2-evoked responses (control vs. MET, P = 0.6338; MET vs. MET + NE, P = 0.9741). n = 7 cells from 4 mice. RM-ANOVA, Sidak’s multiple-comparisons test. Data are mean ± s.e.m. **g**, Recording configuration for calcium currents (I_Ca_) from RBs, and representative traces showing RB I_Ca_ under control and NE conditions (right). **h,i**, Quantification of NE effects on peak amplitude (P = 0.3750) and charge integral (P = 0.4688) of voltage-gated calcium currents recorded from mouse rod bipolar cells. N = 7 cells from 5 mice. ****P < 0.0001; ns, not significant. Paired *t*-test or Wilcoxon test. Unless otherwise indicated, data are presented as individual points with means; error bars are not shown. Related to Fig. 2.

**Extended Data Fig. 9.**
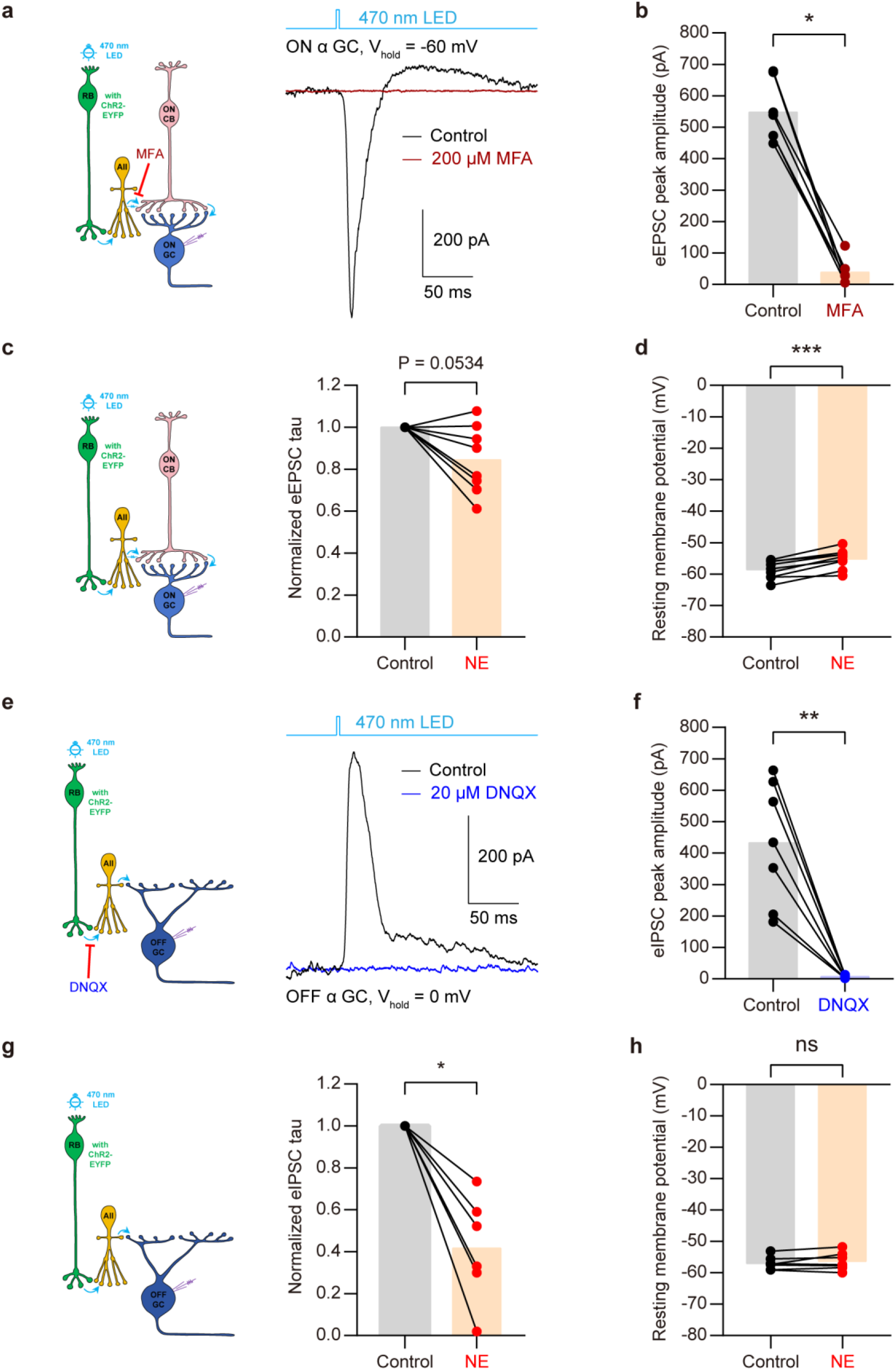
Validation of rod bipolar cell input to α ganglion cells and effects of norepinephrine on response kinetics and membrane potential. **a**, Recording configuration for evoked excitatory postsynaptic currents (eEPSCs) from ON α ganglion cells during optogenetic stimulation of rod bipolar cells (left), and representative eEPSCs showing that meclofenamic acid (MFA; 200 μM), which blocks electrical synapses between AII amacrine cells and ON cone bipolar cells, nearly abolishes the response (right). **b**, Quantification of MFA effect on eEPSC peak amplitude. P = 0.0156. n = 7 cells from 3 mice. **c**, Recording configuration for eEPSCs from ON α ganglion cells (left), and quantification showing that norepinephrine (NE; 100 nM) modestly accelerates eEPSC decay (right). n = 8 cells from 5 mice. **d**, Quantification showing that NE slightly depolarizes ON α ganglion cells. P = 0.0002. n = 9 cells from 5 mice. **e**, Recording configuration for evoked inhibitory postsynaptic currents (eIPSCs) from OFF α ganglion cells during optogenetic stimulation of rod bipolar cells (left), and representative eIPSCs showing that DNQX (20 μM), which blocks AMPA receptors at rod bipolar cell→AII synapses, eliminates the response (right). **f**, Quantification of DNQX effect on eIPSC peak amplitude. P = 0.0011. n = 7 cells from 5 mice. **g**, Recording configuration for eIPSCs from OFF α ganglion cells (left), and quantification showing that NE (100 nM) modestly accelerates eIPSC decay (right). P = 0.0313. n = 6 cells from 4 mice. **h**, Quantification showing that NE does not alter the resting membrane potential of OFF α ganglion cells. P = 0.2940. n = 7 cells from 5 mice. *P < 0.05; **P < 0.01; ***P < 0.001; ns, not significant. Paired *t*-test or Wilcoxon test. Data are presented as individual points with means; error bars are not shown. Related to Fig. 4.

**Extended Data Fig. 10.**
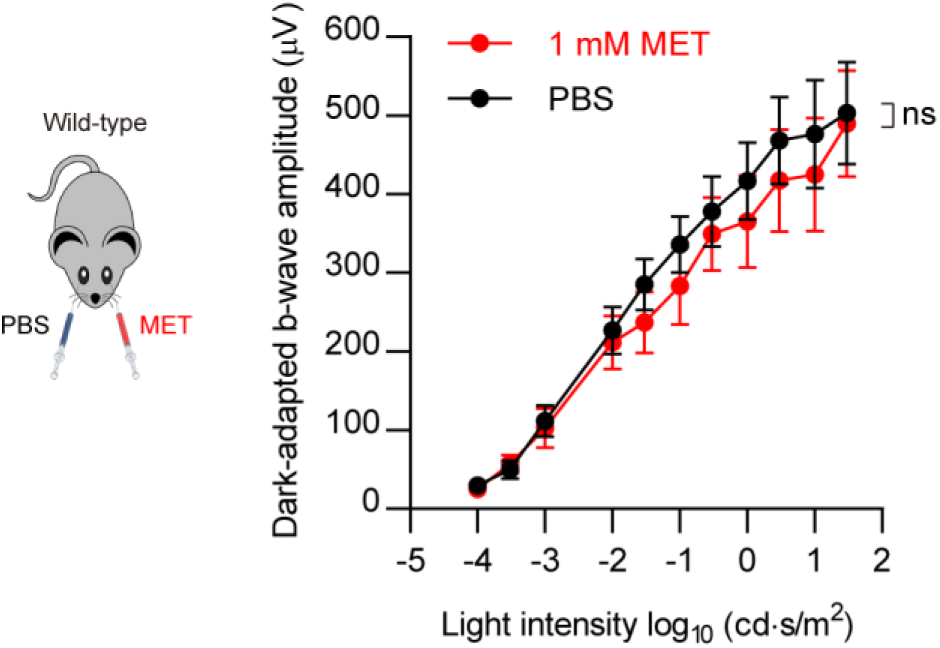
Pharmacological β_1_AR blockade does not alter baseline scotopic electroretinograms. Schematic of scotopic electroretinogram recording after intravitreal injection of metoprolol (MET) or PBS into opposite eyes of the same mouse (left). Intravitreal MET (1 mM) does not significantly alter dark-adapted b-wave amplitudes compared with PBS control across flash intensities (right). ns, not significant. n = 6 mice. Paired *t*-test. Data are mean ± s.e.m. Related to Fig. 5.

## Author contributions

J.B.K. conceptualized the study. F.S.T. and Z.X.W. performed the patch-clamp recording experiments. F.S.T., Y.R.L., and M.G. performed the electroretinogram experiments. M.M.K. and J.B.L. performed the computational analysis, with contribution from J.T.L. L.J.R., Z.X.W., F.S.T., and S.Y.C. performed the single-cell RNA sequencing, fluorescent in situ hybridization, and immunohistochemistry experiments. Y.R.L and T.T.Z bred the *Adrb1* conditional knockout and *Pcp2*-cre; Ai32 mice. Y.C., B.G., C.Y., and H.B.Y. provided human, macaque, and cat tissues, respectively. J.B.K., F.S.T., M.M.K., Y.R.L., Z.X.W., M.G., and J.B.L. analyzed the data. J.B.K., F.S.T., and M.M.K. wrote the manuscript. J.B.K., F.S.T., M.M.K., Y.R.L., Z.X.W., and W.L. edited the manuscript.

## Acknowledgements

This work was supported by start-up funding from Oujiang Laboratory to J.B.K. (OJQD2023003) and by the Fundamental Research Funds of the State Key Laboratory of Ophthalmology, Sun Yat-sen University, to J.B.K. The authors thank Dr. J.H. Singer for helpful discussions; Drs. X.L. Yang, S.J. Weng, Q. Li, and J.H. WU for assistance in obtaining adult human retinal sections; Ms. C.M. Chen for assistance in obtaining Chinese yellow cattle and pig eyes; and the Core Facilities of Oujiang Laboratory for expert technical support. Species icons used in the figures were generated using Doubao. During manuscript preparation, the authors used DeepSeek and ChatGPT to improve readability and language quality. These tools were used solely for language editing and expression refinement, and were not used to generate scientific content, data or analyses. All authors reviewed the final manuscript and take full responsibility for its content.

## Methods

### Ethics and biological materials

#### Human retinal tissue

Human retinal samples were obtained post-mortem from the Eye & ENT Hospital of Fudan University. Specimens were derived from one male donor, aged 83 years, and one female donor, aged 77 years, with no documented history of ocular disease or intraocular surgery. The use of de-identified human retinal tissues was approved by the Ethics Committee of the Eye & ENT Hospital of Fudan University (#2023126). This study was conducted in accordance with the tenets of the Declaration of Helsinki. Informed consent for the tissue donation was obtained from donors or their legal guardians.

#### Non-mouse retinal tissue

Eyes from Chinese yellow cattle and pigs were obtained from an abattoir in Yongjia County, Wenzhou, China, approximately 1 h after death. Additional retinal tissues were obtained from animal colonies maintained at Fudan University (cat; Shanghai, China), Kunming University of Science and Technology (macaque; Kunming, China), Zhejiang Vital River Laboratory Animal Technology Co., Ltd. (rat; Jiaxing, China), and Wenzhou Gaofei Biotechnology Co., Ltd. (rabbit; Wenzhou, China). The collection and use of these tissues were approved by the relevant institutional animal ethics committees. The numbers of animals used for each histological experiment are provided in figure legends.

#### Mice

All procedures involving mice were approved by the Institutional Animal Care and Use Committees of Oujiang Laboratory (#OJLAB25072902) and Sun Yat-sen University (#2018-153) and were performed in accordance with the relevant institutional and national guidelines.

Mice were maintained in a controlled environment with a 12-h light/dark cycle, ambient temperature of 21-23 °C, and relative humidity of 45-55%, with ad libitum access to standard chow and water. Wild-type C57BL/6J mice, channelrhodopsin-2 (ChR2)-expressing mice, and rod-bipolar-cell-specific *Adrb1* conditional knockout mice and their control littermates of both sexes, aged 2-10 months, were used.

Mice expressing ChR2 predominantly in rod bipolar cells were generated by crossing BAC-*Pcp2*-IRES-Cre mice (*Pcp2*-Cre; The Jackson Laboratory, #010536)^51^ with Ai32 mice (The Jackson Laboratory, #012569)^52^. Rod-bipolar-cell-specific *Adrb1* conditional knockout mice were generated by crossing *Pcp2*-Cre mice with *Adrb1*^fl/fl^ mice (GemPharmatech, #T005857). Cre-negative *Adrb1*^fl/fl^ littermates served as controls. Genotypes were determined by PCR with specific primers.

#### Retinal electrophysiology

##### Retinal slice preparation

Vertical retinal slices were prepared from light-adapted adult wild-type C57BL/6J or ChR2-expressing mice as described previously^27,45,53–55^. After euthanasia or decapitation, eyes were rapidly enucleated and retinas were isolated in oxygenated Ames’ medium. Retinas were cut into 200-μm-thick slices using a VT1200S vibratome (Leica Microsystems). Slices were maintained at room temperature in Ames’ medium continuously equilibrated with 95% O₂ and 5% CO₂ until recording.

Recordings were performed at 32-34 °C under continuous superfusion at 1-2 mL/min with oxygenated artificial cerebrospinal fluid (ACSF) containing 119 mM NaCl, 23 mM NaHCO₃, 1.25 mM NaH₂PO₄, 2.5 mM KCl, 1.15 mM CaCl₂, 1.5 mM MgCl₂, 10 mM glucose, 2 mM sodium lactate, and 2 mM sodium pyruvate.

To isolate optogenetically evoked responses arising from rod bipolar cell circuitry while suppressing photoreceptor-driven input to bipolar cells, the ACSF contained L-AP4 (5 μM), a group III metabotropic glutamate receptor agonist that suppresses ON bipolar cell signaling, and ACET (1 μM), a kainite receptor antagonist that suppresses OFF bipolar cell signaling. Where required, GABA_A_-, GABA_C_-, and glycine-receptor-mediated currents were blocked with picrotoxin (100 μM), I4AA (10 μM) or TPMPA (50 μM), and strychnine (0.5 μM), respectively. Tetrodotoxin (TTX; 0.5 μM) was additionally included during miniature and evoked excitatory postsynaptic current recordings from AII amacrine cells.

##### Patch-clamp recording

Patch electrodes were pulled from borosilicate glass capillaries (BF100-50-10, Sutter Instrument). Electrodes used for recordings from rod bipolar cells and AII amacrine cells had resistances of 7-11 MΩ.

For current-clamp recordings, electrodes were filled with an intracellular solution containing 110 mM potassium gluconate, 5 mM NaCl, 10 mM HEPES, 8 mM phosphocreatine, 4 mM Mg-ATP, 0.4 mM Na₃-GTP and 1 mM BAPTA. For voltage-clamp recordings, the intracellular solution contained 95 mM cesium methanesulfonate, 20 mM TEA-Cl, 1 mM 4-aminopyridine, 10 mM HEPES, 8 mM phosphocreatine, 4 mM Mg-ATP, 0.4 mM Na₃-GTP, and 1 mM BAPTA. Solutions were adjusted to pH 7.2 with KOH or CsOH and to approximately 285 mOsm with sucrose.

Cells were targeted using infrared differential interference contrast optics. Alexa Fluor 488 or Alexa Fluor 594 was included in the intracellular solution where required to confirm cellular morphology. Rod bipolar cells and AII amacrine cells were identified on the basis of soma position, morphology, fluorescence and characteristic electrophysiological properties.

During voltage-clamp recordings, rod bipolar cells and AII amacrine cells were typically held at −60 mV and −80 mV, respectively. Membrane potentials were corrected for a liquid-junction potential of approximately −10 mV. Access resistance was less than 40 MΩ for rod bipolar cells and less than 25 MΩ for AII amacrine cells and was compensated by 50-90%. Recordings were excluded if access resistance changed by more than 10% during an experiment.

Signals were amplified using MultiClamp 700B amplifiers (Molecular Devices), sampled at 10-20 kHz and low-pass filtered at 2 kHz. Analogue-to-digital conversion was performed using an ITC-18 interface (HEKA/Instrutech). Data acquisition and stimulation were controlled using programs written in Igor Pro 6 (WaveMetrics).

Miniature events were detected and analyzed using Igor Pro 6. Baseline recordings were acquired over a 3-min epoch, and drug-treatment measurements were obtained 10 min after drug application. The fast and slow components of evoked currents were quantified using the same software.

##### Whole-mount recordings

Retinas from ChR2-expressing mice were dissected in oxygenated Ames’ medium and mounted ganglion-cell-side up on membrane filter paper with an approximately 2-mm-diameter central aperture. Preparations were superfused at 3 mL/min with oxygenated ACSF maintained at 32-34 °C.

Cells with large somata exceeding approximately 20 μm in diameter were targeted in the ganglion cell layer, enriching recordings for ON α and OFF α retinal ganglion cells^27,56^. Cell identity was further assessed from light-response polarity and post hoc morphology after inclusion of Alexa Fluor 594 in the recording pipette.

Whole-cell voltage-clamp recordings were performed using electrodes with resistances of 3-6 MΩ filled with the cesium-based intracellular solution described above, supplemented with QX-314 (5 mM). Current-clamp recordings were performed using the potassium-based intracellular solution.

Excitatory postsynaptic currents were isolated by holding ganglion cells near the chloride equilibrium potential, approximately −60 mV. Inhibitory postsynaptic currents were isolated by holding cells near the reversal potential for ionotropic glutamate-receptor-mediated currents, 0 mV. Access resistance was less than 20 MΩ and was compensated by 50%. The superfusion solution contained L-AP4 (5 μM) and ACET (2 μM) throughout the recordings. Strychnine was omitted when recording glycinergic inhibitory input to OFF α ganglion cells.

Meclofenamic acid (MFA; 200 μM) was applied to block electrical coupling between AII amacrine cells and ON cone bipolar cells. DNQX (20 μM) was used to block AMPA/kainate receptor-mediated transmission at rod bipolar cell→AII amacrine cell synapses.

Peak latency was measured from stimulus onset to the peak synaptic current. First-spike latency was measured from stimulus onset to the first action potential. Electrophysiological data were analyzed using Igor Pro 6.

#### Optogenetic stimulation

ChR2 was activated using a 470-nm high-power LED (Thorlabs) delivered through a 60× objective, which produced a light spot approximately 125 μm in diameter. Light intensity, pulse duration, and inter-stimulus intervals were controlled via Igor Pro 6, as described previously^45,54,55^.

#### Pharmacological agents

Chemicals were obtained from Sigma-Aldrich or Tocris Bioscience unless otherwise stated. Norepinephrine was obtained from Abcam or Tocris Bioscience; NF449 and NF023 from Merck Millipore; SQ22536 from Santa Cruz Biotechnology; and Alexa Fluor-conjugated dyes from Invitrogen.

Stock solutions were prepared in water or DMSO and stored according to the manufacturers’ recommendations. The final DMSO concentration did not exceed 1‰ and was matched in control solutions.

#### Electroretinography (ERG)

Scotopic electroretinographic responses were recorded using a RETI-scan system (Roland Consult). Mice were dark-adapted for at least 8 h, and all subsequent procedures were performed under dim red illumination.

Animals were anaesthetized with tribromoethanol (250 mg/kg, intraperitoneally) based on established protocols^57,58^. To minimize potential adverse effects, including peritoneal adhesions, animals were euthanized immediately following data acquisition. Throughout the experimental period, no anesthesia-related mortality or signs of severe distress were observed. Pupils were dilated using 1% tropicamide and 2.5% phenylephrine, and topical tetracaine hydrochloride was applied for corneal anesthesia. A 1-μL volume of norepinephrine (1 μM), isoproterenol (10 μM), metoprolol (1 mM) or PBS was delivered by intravitreal injection using a 33-gauge Hamilton syringe under a surgical microscope. Drug and vehicle were administered to opposite eyes of the same animal, and injection laterality was randomized across animals.

Animals were maintained at 37 °C throughout the recording. Gold-ring corneal electrodes with a diameter of 3 mm were positioned on both eyes using 1% hypromellose to maintain electrical contact. Reference electrodes were inserted subcutaneously near each eye, and a ground electrode was inserted into the tail.

Recordings began 25 min after injection. Flash intensities ranged from 0.0001 to 30 cd·s/m^2^, with interstimulus intervals of 5-30 s depending on flash intensity. Responses to dim flashes below 0.3 cd·s/m^2^ were averaged over six trials, whereas responses to flashes of 0.3 cd·s/m^2^ or greater were averaged over three trials. The b-wave amplitude was measured from the trough of the a-wave to the peak of the b-wave using the built-in analysis software (Roland Consult).

#### Immunohistochemistry

Adult wild-type mice were euthanized by cervical dislocation, and eyes were rapidly enucleated. After removal of the cornea and lens, eyecups were fixed in 4% paraformaldehyde in PBS for 20 min, washed in PBS and cryoprotected sequentially in 10%, 20% and 30% sucrose. Eyecups were embedded in OCT compound (Sakura Finetek, #4583), sectioned vertically at 14 μm using a Leica cryostat and stored at - 80 °C until staining.

Sections were blocked for 2 h at room temperature in PBS containing 0.1% Triton X-100 and 6% BSA. Primary antibodies were diluted in PBS containing 0.1% Triton X-100 and 3% BSA and applied overnight at 4 °C. The primary antibodies were goat anti-β_1_AR (1:25; Sigma-Aldrich, #SAB2500034) and rabbit anti-PKCα (1:10,000; Sigma-Aldrich, #P4334).

After washing, sections were incubated for 2 h at room temperature in the dark with species-appropriate Alexa Fluor-conjugated secondary antibodies (Invitrogen). For antibody-specificity controls, the β_1_AR antibody was preabsorbed with its immunizing peptide before application to control sections.

Images were acquired using an LSM 880 laser-scanning confocal microscope (Carl Zeiss) equipped with a Plan-Apochromat 63×/1.4 numerical aperture oil-immersion objective. Acquisition settings were kept constant for directly compared samples. Brightness and contrast were adjusted uniformly across images using ZEN (Carl Zeiss) and Adobe Photoshop. No selective or nonlinear image manipulation was applied.

#### RNAscope in situ hybridization

RNAscope multiplex fluorescent in situ hybridization was performed on retinal cryosections from wild-type and *Adrb1* conditional knockout mice, rats, cats, pigs, cattle, macaques, and humans using the RNAscope Multiplex Fluorescent Reagent Kit v2 with the integrated co-detection workflow (Advanced Cell Diagnostics, #323100).

Eyes were fixed in 4% paraformaldehyde at 4 °C for 2 h or overnight, depending on species. Retinas were dissected, cryoprotected sequentially in 10%, 20%, and 30% sucrose, embedded in OCT compound and sectioned at 14 μm.

Sections underwent target retrieval and protease treatment before hybridization with species-specific probes against *ADRB1/Adrb1*, *Adra2a*, *Adra2b* or *Adra2c*. All the probes were purchased from Advanced Cell Diagnostics. Mouse probe catalogue numbers included *Adrb1* (449761), *Adra2a* (425341), *Adra2b* (425321) and *Adra2c* (846271). Human probe catalogue number included *ADRB1* (469511). Fluorescent signals were developed according to the manufacturer’s protocol.

For immunofluorescence co-detection, sections were washed in PBS and blocked for 2 h at room temperature with 6% normal donkey serum. Sections were incubated overnight at 4 °C with one or more of the following antibodies diluted in 3% normal donkey serum: rabbit anti-PKCα (1:5,000; Sigma-Aldrich, #P4334), rabbit anti-glycine transporter 1 (1:500; Synaptic Systems, #272103) and mouse anti-parvalbumin (PV; 1:500; Swant, #PV235PUR).

Species-appropriate Alexa Fluor-conjugated secondary antibodies were applied for 2 h at room temperature. Sections were counterstained with DAPI, mounted and imaged using an LSM 980 confocal microscope (Carl Zeiss) with 20× or 63× objectives. Samples being directly compared were acquired using identical imaging settings.

#### Cross-species single-cell analysis

##### Dataset processing

All single-cell RNA-seq datasets analyses were performed in R using Seurat v5.3.1, developed by the Satija laboratory (https://satijalab.org/seurat/), together with Matrix v1.7.2, devtools v2.4.5, dplyr v1.1.4, and ggplot2 v4.0.1 for data processing, statistical analysis and visualization.

Each species data was respectively reanalyzed using a uniform processing and normalization workflow. Species identities were added to the metadata, and species-specific objects were merged for comparative visualization. Expression comparisons were performed using normalized expression values from the RNA assay.

##### Retinal cell annotation

Each cluster was assigned to a retinal cell type based on the expression of orthologues of canonical markers characterized in the mouse retina. For consistency across species, orthologous gene symbols were standardized to uppercase. The markers used for annotation were as follows: rods (*RHO*, *PDE6A*, and *NRL*), cones (*ARR3*, *PDE6H*, and *OPN1MW*), horizontal cells (*LHX1*, *TPM3*, and *ONECUT1*), rod bipolar cells (*VSX1*, *OTX2*, and *PRKCA*), ON cone bipolar cells (*VSX1*, *OTX2*, and *GRM6*), OFF cone bipolar cells (*VSX1*, *OTX2*, and *GRIK1*), glycinergic amacrine cells (*SLC6A9* and *TFAP2A*), GABAergic amacrine cells (*GAD1* and *GAD2*), retinal ganglion cells (*RBPMS*, *THY1*, and *SLC17A6*), Müller glia (*RLBP1*, *AQP4*, and *APOE*), microglia (*CX3CR1*, *CSF1R*, and *AIF1*), and astrocytes (*GFAP*, *PAX8*, and *S100B*).

Cell-type assignments were based on combinations of markers rather than the expression of any single gene. Ambiguous or low-quality clusters were excluded and not used in subsequent analysis.

#### Rabbit single-cell RNA sequencing

##### Tissue dissociation

Retinas were collected from two adult male rabbits. Retinal tissue was dissected and minced using a sterile blade, then transferred into a 15 mL tube containing tissue dissociation medium supplemented with Liberase^TM^ (Roche, #05401127001) and DNase I (Solarbio, #D8071), and incubated for 50 min at 37 °C.

Dissociated cells were filtered through a 70-μm cell strainer and centrifuged at 300 g for 7 min at 4 °C. Cell pellets were resuspended in D-PBS supplemented with 10% BSA and subjected to AO/PI staining to assess cell concentration and viability. Dead cells were removed using a Miltenyi Biotec Dead Cell Removal Kit (Miltenyi Biotec, #130-090-101) according to the manufacturer’s protocol. Following washing, the final cell suspension was used immediately for library construction.

##### Library preparation and sequencing

Single-cell libraries were prepared using the Chromium Single cell 3’ Library and Gel Bead Kit V3.1 (10x Genomics, #PN1000268). Cell suspensions were loaded onto the Chromium platform to encapsulate individual cells with barcoded gel beads in emulsions (GEM). Within each emulsion, polyadenylated transcripts were captured and reverse-transcribed into cell-barcoded, UMI-labelled cDNA. After emulsion breakage, cDNA was recovered, amplified, purified and used for next-generation sequencing library construction. Libraries were sequenced on an Illumina NovaSeq 6000 using 150-bp paired-end reads (Berry Genomics Corporation, Beijing, China).

##### Data processing and analysis of single-cell RNA sequencing data

Sequencing reads were processed using Cell Ranger v7.1.0 (10x Genomics) and aligned to the rabbit reference genome (mOryCun1.1). Gene-barcode count matrices were generated using the default Cell Ranger pipeline.

The two rabbit samples were processed independently before integration. Doublets were identified using Scrublet and doublet cells scores greater than 0.25 were classified as putative doublets and removed. Genes detected in > 3 cells, cells with > 200 detected genes, > 400 unique molecular identifier counts and < 20% mitochondrial gene content were retained. After quality control, 14,466 and 12,914 cells were retained from the two samples, respectively, yielding 27,380 cells in total. Each sample was normalized and 2000 highly variable genes were identified. Integration anchors were identified using FindIntegrationAnchors, and the samples were integrated using IntegrateData. The integrated assay was used for dimensionality reduction and clustering, whereas gene-expression visualization was performed using the normalized RNA assay.

The integrated data was then used for scaling, principal-component analysis (PCA), nearest-neighbor graph construction, Louvain clustering and Uniform Manifold Approximation and Projection (UMAP) visualization. Cell types were manually annotated based on cluster-enriched genes, known retinal markers and published literature.

#### Statistics and reproducibility

No statistical method was used to predetermine sample size. Sample sizes were selected on the basis of previous retinal electrophysiology and ERG studies and are reported in the figure legends. Unless otherwise specified, n denotes individual cells for electrophysiological experiments and individual mice for ERG experiments. Electrophysiological recordings for each condition were obtained from at least three independent animals.

Both male and female mice were included. The study was not specifically powered to detect sex-dependent effects, and data from both sexes were pooled. Injection laterality in paired-eye ERG experiments was randomized. Experimental allocation, data acquisition, and analysis were not performed blinded. No data were excluded except on the basis of prespecified technical criteria. Each histological result was reproduced in at least two biologically independent samples.

Statistical analyses were performed using GraphPad Prism v8.3.0. Normality was assessed using the Shapiro-Wilk test. Comparisons between two independent groups were performed using two-sided unpaired Student’s *t*-tests for normally distributed data or Mann-Whitney U-tests for non-normally distributed data. Paired observations were analyzed using two-sided paired Student’s *t*-tests or Wilcoxon matched-pairs signed-rank tests, as appropriate.

Repeated-measures datasets were analyzed using repeated-measures analysis of variance followed by Šidák’s multiple-comparisons test, or using the Friedman test when parametric assumptions were not met. All statistical tests were two-sided unless otherwise stated. P < 0.05 was considered statistically significant. Exact P values, statistical tests and sample sizes are reported in the figure legends or Source Data. Unless otherwise stated, data are presented as mean ± s.e.m.

#### Code availability

Custom Igor Pro programs, kindly provided by the Jeffrey Diamond Laboratory at the NIH, were used in our experiments for stimulus control, electrophysiological recording, and data analysis. All single-cell RNA sequencing analyses were performed using standard software packages as described in the Methods. No new computational algorithms were developed for this study.

#### Data availability

The raw and processed rabbit retinal single-cell RNA-sequencing data generated in this study are available through the Gene Expression Omnibus (GEO) under accession GSE336783. Previously published retinal datasets were obtained from GEO under accession numbers GSE237206 (marmoset), GSE237210 (rhabdomys), GSE237208 (peromyscus), GSE237212 (squirrel), GSE237211 (sheep), GSE237202 (cow), GSE237209 (pig), GSE237203 (ferret), GSE237207 (opossum), and GSE237213 (tree shrew)^5^; GSE118480 (macaque)^35^; GSE63472 (mouse)^34^; and GSE149715 (mouse amacrine cells)^47^. Human retinal datasets were obtained from the European Genome-phenome Archive under accession number EGAS00001004561^36^. Access to controlled human data is subject to the EGA data-access procedures.

